# Fine-tuning molecular mechanics force fields to experimental free energy measurements

**DOI:** 10.1101/2025.01.06.631610

**Authors:** Dominic Rufa, Joshua Fass, John D. Chodera

**Affiliations:** Computational and Systems Biology Program, Sloan Kettering Institute, Memorial Sloan Kettering Cancer Center, New York, NY 10065; Tri-Institutional PhD Program in Chemical Biology, Weill Cornell Graduate School of Medical Sciences, New York, NY 10065, USA; Computation, Relay Therapeutics, Cambridge, Massachusetts 02139, United States

**Author notes:** **For correspondence:** (JDC).

## Abstract

Alchemical free energy methods using molecular mechanics (MM) force fields are essential tools for predicting thermodynamic properties of small molecules, especially via free energy calculations that can estimate quantities relevant for drug discovery such as affinities, selectivities, the impact of target mutations, and ADMET properties. While traditional MM forcefields rely on hand-crafted, discrete atom types and parameters, modern approaches based on graph neural networks (GNNs) learn continuous embedding vectors that represent chemical environments from which MM parameters can be generated. Excitingly, GNN parameterization approaches provide a fully end-to-end differentiable model that offers the possibility of systematically improving these models using experimental data. In this study, we treat a pretrained GNN force field—here, espaloma-0.3.2—as a *foundation simulation model* and *fine-tune* its charge model using limited quantities of experimental hydration free energy data, with the goal of assessing the degree to which this can systematically improve the prediction of other related free energies. We demonstrate that a highly efficient “one-shot fine-tuning” method using an exponential (Zwanzig) reweighting free energy estimator can improve prediction accuracy without the need to resimulate molecular configurations. To achieve this “one-shot” improvement, we demonstrate the importance of using effective sample size (ESS) regularization strategies to retain good overlap between initial and fine-tuned force fields. Moreover, we show that leveraging low-rank projections of embedding vectors can achieve comparable accuracy improvements as higher-dimensional approaches in a variety of data-size regimes. Our results demonstrate that linearly-perturbative fine-tuning of foundation model electrostatic parameters to limited experimental data offers a cost-effective strategy that achieves state-of-the-art performance in predicting hydration free energies on the FreeSolv dataset.

## 1 Introduction

Computational molecular property prediction has become an indispensable resource in drug discovery, offering a means to evaluate critical properties of candidate compounds—such as solubility [89], permeability [63, 104], binding affinity [74], target selectivity [2], and mutational resistance [43, 96]—long before synthesis and experimental assays. While these computational predictions often lack the accuracy of experimental measurements, their speed and cost-efficiency make them highly advantageous [71]. A computational evaluation can be completed in hours without the need to synthesize the compound, compared to the days to months required to synthesize compounds and perform laboratory assays. This rapid turnaround enables researchers to screen and prioritize designs—potentially including large libraries of molecules—and focus resources on the most promising candidates, thereby accelerating the drug discovery process [87].

Among the diverse computational approaches available, including data-driven methods such as machine learning models and cheminformatics, physics-based methods stand out for their versatility and rigor. Rooted in statistical thermodynamics, these methods provide a theoretical grounding for computational molecular property prediction [17]. Many relevant properties—such as hydration free energies, partition coefficients, binding affinities, selectivities, and the impact of mutations—can be directly cast as thermodynamic quantities. These thermodynamic properties are accessible through alchemical free energy calculations that utilize molecular dynamics simulations [40], which serve as a convenient method to sample the equilibrium conformational distribution of the potential energy surface [95], a consequence of the underlying model being used. Low-level models, such as molecular mechanics, approximate the potential energy surface with classical force fields, while higher-level models, such as density functional theory (DFT) [33], provide a more detailed quantum mechanical representation [38, 86]. By preserving the underlying physics and offering a consistent framework for diverse property predictions, physics-based methods uniquely enable robust and mechanistic insights into molecular behavior in addition to quantitative predictions of experimentally measurable properties.

There is an inherent trade-off between speed and accuracy when choosing computational methods for molecular dynamics simulations or other physics-based molecular property prediction approaches. Higher levels of theory, such as density functional theory (DFT) and coupled cluster methods (e.g., CCSD(T)) [5], provide highly accurate quantum mechanical descriptions of molecular interactions, including complex many-body effects. However, these methods are computationally expensive, often limiting their applicability to small systems or short simulation times [14]. Hybrid methods, such as QM/MM (Quantum Mechanics/-Molecular Mechanics), combine quantum mechanical accuracy with the efficiency of molecular mechanics but are still limited by their computational cost, making them impractical for large-scale applications [35, 62]. Hybrid ML/MM (Machine Learning/Molecular Mechanics) approaches [32, 85, 91] and ML (Machine Learning) potentials [4, 6, 55, 79, 98, 106] also offer promising alternatives, as they can capture more detailed interactions compared to classical MM, but they come with their own challenges, including computational efficiency and the need to demonstrate that they are more competitive than state-of-the-art MM methods. In contrast, molecular mechanics (MM) force fields [20, 39, 68], which are time-tested and well-studied, offer a much more computationally efficient alternative, recovering many thermodynamic quantities with orders of magnitude less compute than QM methods due to their simplified functional forms (e.g., Coulombic interactions, low-order Taylor series expansions for bond and angle deviations, simple Fourier expansions for periodic torsions, and Lennard-Jones potentials), the ease of parallelization (e.g., using neighbor lists, unit cells, domain decomposition, and particle decomposition), GPU acceleration (e.g., OpenMM [25], Gromacs [107], NAMD [78], AMBER [13], timemachine [29]), and the comparatively low complexity of software implementation, thanks to the diversity and accessibility of relevant open-source Pythonic libraries.

Despite their classical approximations, MM force fields have proven capable of accurately predicting key molecular and molecular interaction thermodynamic properties, such as those relevant to ADMET and potency (e.g., protein-ligand binding free energies, etc.), often within an accuracy of 1–2 kcal mol^−1^ [18, 93]. This level of accuracy is still capable of significantly accelerating preclinical drug discovery [93], making MM force fields a practical and effective choice for large-scale molecular screening and prioritization in drug discovery campaigns [88].

Even modest improvements in binding affinity prediction accuracy can significantly increase the likelihood of prioritizing more potent compounds for synthesis, thereby reducing time and costs for lead optimization [93]. Motivated by the prospective benefits of increased accuracy, many efforts have been directed towards improving the reliability of MM force field parameterizations to better reflect the energetics and quantum mechanical properties of diverse datasets of biological and organic molecules [10, 45, 69]. The expectation is that achieving a higher correlation between MM and QM energetics will improve the accuracy of molecular property predictions when compared to experimental data [101].

The prospect of improved accuracy through data-driven approaches has driven the development of neural MM force field parameterization approaches, such as Espaloma [100, 111] and other graph neural network-based models [16, 36, 92, 112]. These engines are enabled by the availability of efficient, opensource, differentiable optimization libraries (e.g., PyTorch [3], JAX [12]) that support GPU acceleration, as well as the growing availability of quantum mechanical datasets (e.g., QM9 [81], SPICE [24, 26]), which cover a wide range of chemical space relevant to drug discovery.

Neural MM force field parameterization methodologies are particularly appealing for several reasons. First, they provide a unified and generalizable approach to parameterizing diverse chemical spaces relevant to drug discovery [111]. This overcomes the limitations of legacy methods, which required different force fields for various types of biological molecules, such as carbohydrates (e.g., GROMOS [90]), lipids (e.g., Slipids [49]), and proteins (e.g., OPLS-AA [52] or CHARMM [116]). These inconsistencies led to challenges in integrating different parameterizations into a single molecular dynamics engine, complicating simulations and making it harder to ensure compatibility across different systems [68].

Furthermore, neural MM force field parameterization models eliminate the need for hand-crafted, expert-guided discrete atom-type schemes typically used in legacy MM force fields. Instead, they employ continuous embedding vectors to encode the local chemical environment of each atom. This data-driven approach offers greater robustness, capturing subtle variations in chemical environments more effectively than traditional methods, thereby reducing reliance on potentially fallible domain expertise and intuition [111].

The espaloma model, in particular, initially saw remarkable success in its ability to fit MM force field parameters to reproduce QM energies on a variety of datasets [111]. Its consistent ability to achieve state-of-the-art accuracy on matching QM energies in a subsequent study involving the more diverse SPICE [24, 26] dataset demonstrated its robustness across chemical spaces. This success motivated an investigation into the model’s ability to recover biomolecular thermodynamic properties of condensed-phase matter, as it applies to drug discovery, including protein-ligand binding affinity predictions and NMR spectra of peptides [100]. Interestingly, while espaloma-0.3.2 [100] achieved competitive accuracy in comparison with legacy force fields, namely ff14SB [64]/openff-2.1.0 [10], its inability to achieve a statistically significant improvement in small molecule binding free energies [100] highlighted an important point: improvements in QM energy/force matching alone do not necessarily translate to improvements in accuracy with respect to the relevant properties implicated drug discovery. This suggests that simply improving QM potential energy fits may not be sufficient for enhancing practical predictive power in drug discovery contexts. It is an open question as to whether the inability of MM potentials like espaloma-0.3.2 with improved accuracy to gasphase QM potentials are an inherent limitation of the MM functional form, an inability to accurately model many-body QM interactions, or a failure in modeling the *effective potential* needed to accurately capture omitted effects like quantum nuclear effects, which condensed-phase properties may provide information about.

Nevertheless, accurate MM models built from large amounts of quantum chemical data such as espaloma-0.3.2 are poised as a highly effective tool for enhancing the predictive power of MM-based modeling in pharmaceutical research. Complementing the initial bottom-up, QM-fitting approach of espaloma with an empirical, top-down fine-tuning with experimental data holds the potential to significantly enhance its accuracy. Indeed, refitting potentials to experimental data via automatic differentiation of physical models [31] has already seen success in applications to a coarse-grained water model [102] and even implicit solvent-based hydration free energy predictions [83]. Fine-tuning to experimental data is particularly important for addressing the aforementioned limitations of quantum chemical data, which often fail to capture critical interactions associated with condensed-phase thermodynamic properties [66]. Incorporating experimental data like binding affinity and hydration free energy data into the training pipeline could enable espaloma to achieve state-of-the-art performance in predicting experimental observables that are essential for preclinical drug discovery [11, 15]. This key insight forms the foundation for our subsequent investigation.

## 2 Pretrained espaloma-0.3.2 “foundation” model serves as a baseline standard for accuracy on the FreeSolv hydration free energy dataset, providing a basis for potential enhancement through low-rank electrostatic fine-tuning

Here, we detail the methodology of the espaloma-0.3.2 foundation model and its performance in predicting small-molecule hydration free energies from the FreeSolv [72] dataset using alchemical absolute hydration free energy calculations. We hypothesize that these results suggest that fine-tuning espaloma with a subset of experimental data could lead to enhanced prediction accuracy for hydration free energies.

### 2.1 The espaloma graph neural network (GNN) generates continuous atom embeddings and MM parameters

The espaloma foundation model operates in three stages [111] (see **Figure 1**, *left*):

**Figure 1.**
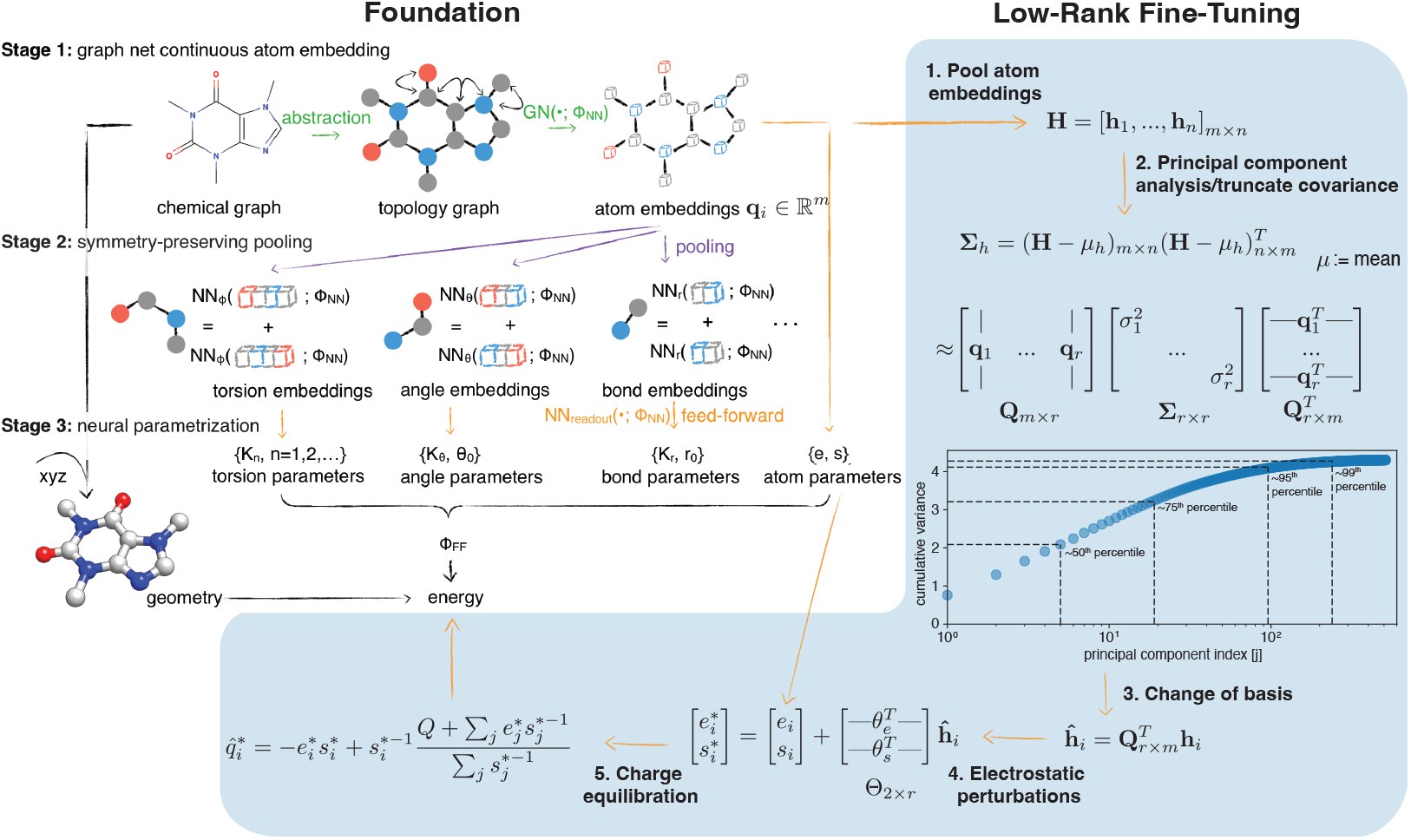
Our approach to rapidly fine-tune an existing espaloma “foundation model” employs a a fast, low-rank perturbation to electrostatic parameters. An existing espaloma graph model for MM parameters (here, espaloma-0.3.2), typically trained with large quantum chemical datasets of molecular energies and/or forces, provides an excellent foundation for fine-tuning to further improve its accuracy on a specific or heterogeneous dataset, possibly of experimental data. We adopt a rapid fine-tuning approach that uses a low-rank approximation to perturb the electrostatics component of the model without needing to re-optimize all model parameters. Fine-tuning proceeds in sequential steps: **1. Pool atom embeddings**: the dataset’s embedding vectors **h**_*i*_ from the foundation espaloma model are pooled into a data matrix **H. 2. Principal component analysis/truncate covariance**: the covariance matrix Σ_*h*_ of the data matrix **H** is diagonalized and the first r principal component vectors are extracted as a truncated matrix **Q**_*m*×*r*_ of orthonormal principal components. The cumulative variance as a function of the number of principal components is shown as an insert. **3. Change of basis**: each **h**_**i**_ is projected into the lower dimensional basis **Q**_*m*×*r*_, yielding 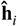. **4. Electrostatic perturbations**: for each atom, the electrostatic parameter vector (*e, s*)^*T*^ representing electronegativity and hardness, respectively, is perturbed by projecting 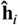 onto two parameterized vectors *θ*_*e*_ and *θ*_*s*_. **5. Charge equilibration**: the new atomic partial charges 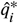 for each atom is recomputed using Charge equilibration (QEq).

### 1. Graph neural network embeddings

the espaloma GNN performs graph convolutions via message passing, which annotates each atom with continuous atom embeddings **h**_*i*_, replacing rule-based atom typing and preserving chemical symmetries. Each **h**_*i*_ encodes molecular connectivity, hybridization, element, and other chemical environment details into a single 512-dimensional real-valued vector.

### 2. Symmetry-preserving pooling

Subsequent embeddings for 2-, 3-, and 4-body terms are evaluated by combining the **h**_*i*_ vectors in a nonlinear, symmetry-preserving (i.e., reflection-invariant) fashion. Notably, and a key focus of our investigation, one-body **h**_*i*_ terms are bypassed directly to Stage 3.

### 3. Neural parameter assignment

The 1- 2-, 3-, and 4-body embeddings from stage 2 are passed to a nonlinear neural network readout function (parameterized by Φ_*NN*_) to generate charge (electronegativity parameters e, hardness parameters s), bond, angle, and torsion parameters, respectively.

Subsequently, the MM parameters and user-provided atomic position vectors are passed to a molecular dynamics engine (timemachine [29], in our investigation) with a differentiable energy function to evaluate energies and forces with respect to atomic position vectors. In this study, we specifically extract the potential energies of trajectories to compute absolute hydration free energies.

To develop a broadly applicable, self-consistent MM forcefield for biomolecular modeling, a high-quality gas-phase quantum chemical dataset was curated from QCArchive [97] in a previous publication [100]. This dataset includes data from multiple sources, providing comprehensive coverage of relevant biomolecular chemistries. The components include: SPICE [24], OpenFF 1.x (“Parsley”), 2.x (“Sage”), and 3.x (“Rosemary”) [108], PepConf [80]. In total, the datasets consist of over 1.18 million conformations across 17,427 unique molecules [100]. AM1-BCC ELF10 [47, 48] partial charges were computed using the OpenEye Toolkits [75] to train Espaloma on AM1-BCC quality charges. Quantum chemical energies were calculated using the Open Force Field standard level of theory (B3LYP-D3BJ/DZVP [37]) with the Psi4 [105] quantum chemistry package.

### 2.2 FreeSolv serves as a valuable dataset with which to assess the accuracy of the espaloma-0.3.2 foundation model on hydration free energy predictions

The FreeSolv dataset [72] is a curated collection of 642 neutral, organic molecules, many of which have drug-like properties or moieties, making them highly relevant to preclinical drug discovery campaigns. It is a critical resource for evaluating and refining MM force fields, particularly for hydration free energy calculations. By providing a curated set of experimental hydration free energies for small neutral molecules in water, FreeSolv enables systematic benchmarking of computational methods, allowing researchers to assess the accuracy and limitations of force fields in capturing solvation phenomena [21, 28, 41, 53, 82, 115]. Such benchmarking is essential for identifying discrepancies between calculated and experimental data, which in turn informs the development of more robust and predictive force fields.

Previous studies using various open-source force fields have demonstrated accuracies in hydration free energy predictions ranging from approximately 2.1 to 1.5 kcal mol^−1^ [22, 23]. Given FreeSolv’s diverse chemical space and ample number of data points, we postulated that this dataset would be a strong candidate for fine-tuning espaloma-0.3.2, as it allows for meaningful partitioning into training and testing sets. The dataset’s comprehensiveness—offering annotated molecular structures and input files—facilitates reproducibility and cross-method comparisons, making it particularly valuable for assessing the accuracy of solute-solvent interactions encoded in molecular mechanics models. Furthermore, the emphasis on hydration free energies—a property that is both experimentally accessible but computationally challenging to accurately predict—underscores FreeSolv’s utility for evaluating the predictive accuracy of solvation models. The diverse set of molecules in FreeSolv also provides a statistically meaningful basis for assessing the generalizability of force fields across a wide range of chemical environments, and enables the iterative refinement of computational models.

### 2.3 Absolute hydration free energy calculations are performed with an alchemical 4D decoupling strategy

Furthermore, we pursue absolute hydration free energy predictions in this study because of their relative ease of implementation and low computational effort to perform highly precise calculations as compared to more complex systems (i.e., protein-ligand relative binding free energy calculations). In our approach, a one-dimensional alchemical protocol is defined, transitioning from *λ* = 0 → 1. At *λ* = 0, the nonbonded (i.e., electrostatic and steric) interactions of the small molecule are “decoupled” from the TIP3P [65] solvent with a 4D lifting strategy [84]. As *λ* increments from 0 to 1, a predefined protocol gradually shrinks the 4D projection, and hence, the corresponding nonbonded interactions, from 1.2Åto 0 (fully coupled) at *λ* = 1. More explicitly, the radial “lifting” is applied to each alchemical (i.e., small-molecule atom) nonbonded term (Lennard-Jones [60] and electrostatic, respectively) through the radial term via

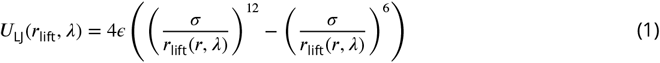

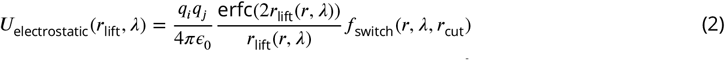

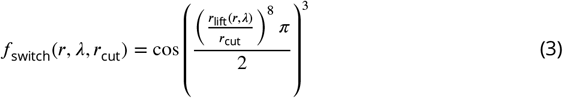

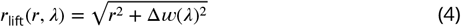

where *q*_*i*_ and *q*_*j*_ are the charges of two particles, *r* is their Euclidean distance between two particles *i* and *j* in ℝ^3^, and Δ*w*(*λ*) is the difference between the corresponding particles’ fourth dimensional lifting coordinate *w*(*λ*). *w*(*λ*) is a pre-defined linear spline in *λ* (see **Supplement**). The electrostatic potential energy *U*_electrostatic_ includes exclusively the direct space coulombic interaction with a multiplicative switching function *f*_switch_(*r, λ, r*_cut_) that decays smoothly to zero at *r*_cut_ (see Eq. 3). The reciprocal space contribution is omitted, as is convention with the timemachine library. *r*_cut_ is 12Å, and interactions beyond *r*_cut_ are omitted. Finally, only molecule-solvent interactions are lifted in the alchemical protocol, preserving the full steric and electrostatic energetics of solvent-solvent and molecule-molecule interactions.

### 2.4 The espaloma-0.3.2 foundation model recovers adequate correlations with small-molecule hydration free energies from the FreeSolv dataset with potential for improvement via electrostatic fine-tuning

Espaloma 0.3.2 [100] / TIP3P [50] demonstrates promising accuracy in hydration free energy predictions, shown in **Figure 2**, though there remains significant potential for improvement. We hypothesize that the observed errors in hydration free energy predictions stem from inaccurate modeling of the appropriate amount of molecular polarization as molecules transition from vacuum to water [76, 84, 99, 99]. Since espaloma-0.3.2 was primarily trained on vacuum-phase quantum chemical (DFT) energies and AM1-BCC partial charges, it likely does not fully capture condensed-phase polarization effects that manifest in modified partial atomic charges, contributing to the large observed root mean square error (RMSE). Additionally, traditional MM force fields do not explicitly include many-body intermolecular interactions, likely causing further deviations from experimental hydration free energies.

**Figure 2.**
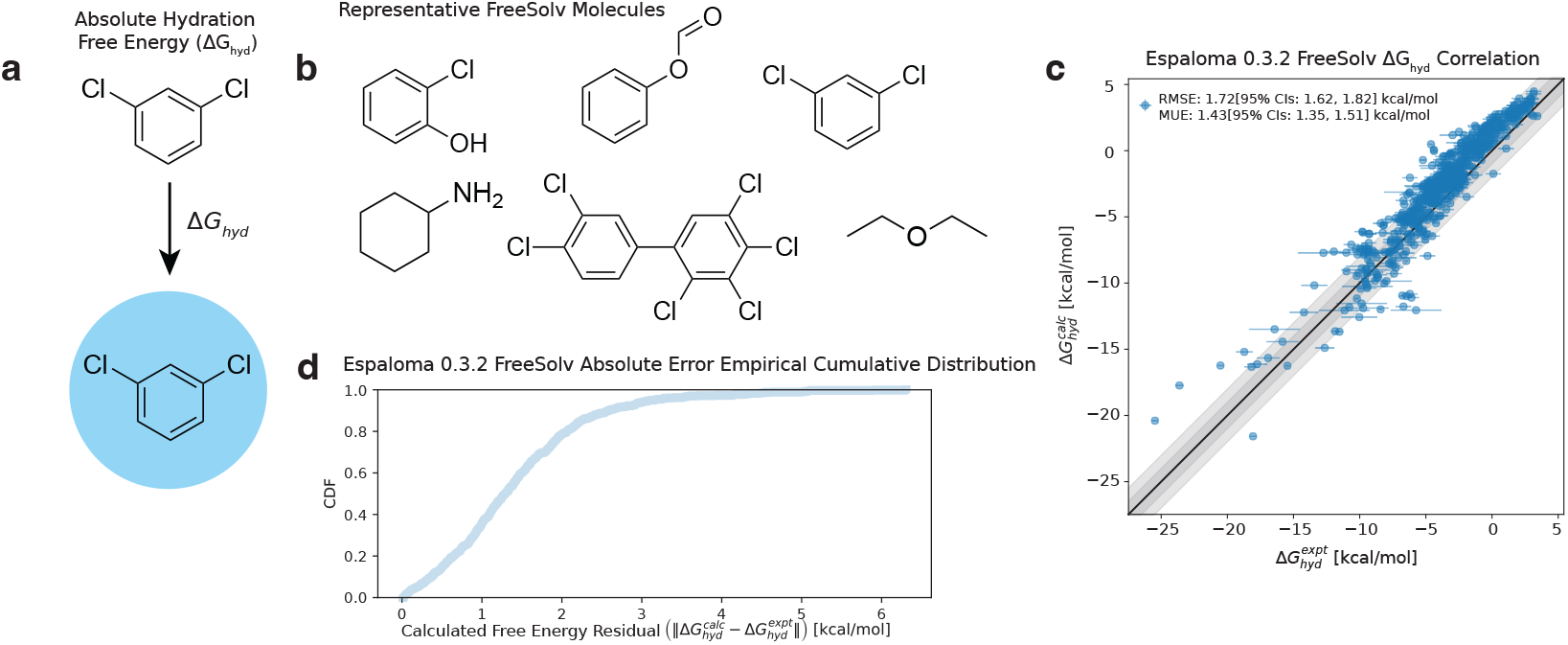
The espaloma-0.3.2 foundation model gives high correlation with experiment on the FreeSolv small molecule hydration free energy benchmark, but leaves room for systematic improvement. **(A)** Absolute hydration free energy calculations transport small molecules from vacuum to water by alchemically coupling intermolecular nonbonded interactions between small molecules and the water solvent. **(B)** Six representative molecules in the Free-Solv dataset, which consists of 642 neutral small organic molecules. **(C)** The espaloma-0.3.2 foundation model’s absolute hydration free energy calculations correlate well with experimental hydration free energies. **(D)** Empirical cumulative distribution function of absolute residuals between calculated and experimental hydration free energies.

To mitigate the errors linked to poorly modeled polarization specifically, we propose fine-tuning the espaloma atomic partial charge model [100, 110, 112] by modifying the electronegativity (*e*) and hardness (*s*) parameters based on a representative subset of the FreeSolv dataset [23]’s experimental hydration free energies. We posit that empirically fine-tuning hydration free energies on a subset of experimental data with representative coverage of chemical space across the full FreeSolv dataset [23] will afford improvements in free energy predictions across the rest of the dataset. Fine-tuning the partial charge model is relatively straightforward compared to adjusting steric parameters or many-body interactions, which are more challenging to modify (see **Discussion**). This approach of charge fine-tuning, as opposed to more intricate empirical fine-tuning of steric or many-body terms, can be incorporated into the model more readily. A detailed discussion of these approaches, particularly the comparison between empirical fine-tuning techniques and first-principles methods, will be presented in **Discussion**.

## 3 Low-rank electrostatic fine-tuning via Zwanzig reweighting and Charge Equilibration (QEq) is an efficient route to fine-tune the espaloma-0.3.2 foundation model to experimental data

### 3.1 Fine-tuning uses atom embeddings to derive data-driven, low-rank perturbations to electrostatic parameters, which modify atomic partial charges via Charge Equilibration (QEq)

In line with our hypothesis that modifying atomic partial charges can enhance the accuracy of hydration free energy predictions, we adopt a data-driven fine-tuning approach which leverages the intermediate parameters generated by the foundation model, namely existing electrostatic parameters and atom embedding vectors. This fine-tuning process, outlined as the “Fine-Tuning” step highlighted in blue in **Figure 1**, proceeds as follows:

#### 1. Pool atom embeddings

The atom embedding vectors **h**_*i*_ ∈ ℝ^*m*^ (*m* = 512) generated by espaloma-0.3.2 are concatenated as separate columns into a data matrix **H**_*m*×*n*_ = [**h**_1_, …, **h**_*n*_] _*m*×*n*′_ implying *n* atoms total in the full FreeSolv dataset.

#### 2. Principal component analysis/truncate covariance

In order to find the *r* orthogonal directions of highest variance in the dataset’s atom embeddings, principal component analysis (PCA) is used to diagonalize a data-derived covariance matrix, which is truncated at *r* principal component vectors as follows:

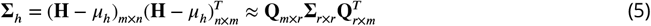

where Σ_*h*_ is the aforementioned covariance matrix, *μ*_*h*_ is the mean embedding vector, **Q**_*m*×*r*_ is the first *r* orthogonal principal component vectors by column, and Σ_*r*×*r*_ is the diagonal, truncated variance matrix with corresponding variances 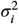 in descending order.

#### 3. Change of basis

Each atom embedding vector **h**_*i*_ ∈ ℝ^*m*^ is projected onto the low-rank basis set of principal component vectors **Q**_*m*×*r*_ to recover the most distinguishing (maximally-variant) embedding features. The low-rank embedding vector is given by

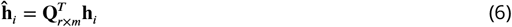

#### 4. Electrostatic perturbations

The low-rank embedding vectors 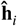 are linearly transformed with a fine-tuning model parameter matrix **Θ**_2×*r*_ to compute perturbations to the foundation model’s electrostatic parameters (*e, s*)^*T*^, (electronegativity and hardness, respectively). The full perturbation is given by

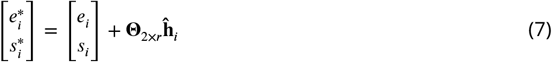

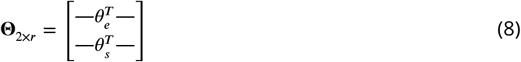

where *θ*_*e*_ and *θ*_*s*_ are learnable parameter vectors that perturb *e* and *s*, respectively.

#### 5. Charge equilibration

Finally, the perturbed electrostatic parameters are passed to the charge equilibration (QEq) [34] equation,

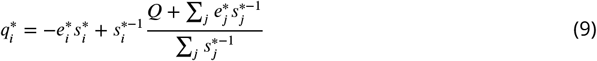

which computes modified atomic partial charges 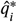 and ensuring that the total molecular charge *Q* remains unperturbed. The relationship between fine-tuning model parameters **Θ** and 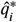 can be made more explicitly with

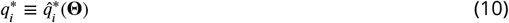

where the dependence on *e, s*, **ĥ**_*i*_, and *q*_*i*_ have been suppressed for clarity. Also, because of the linearity of the *e* and *s* perturbations given by Eq. 7,

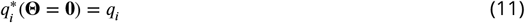

where *q*_*i*_ is the original espaloma-0.3.2 partial charge.

As a further clarification, the QEq equation defines the electronegativity and hardness as the first and second derivatives of a fictitious (and conformation-independent) electostatic potential energy with respect to atomic charge [112]. It represents a second-order Taylor series expansion of the electrostatic energy with a Lagrange multiplier used to constrain the total molecular charge *Q*.

This fine-tuning mechanism is motivated by several key insights. First, we apply PCA to project the existing embedding vectors **h**_*i*_ into a low-rank space, serving the dual purpose of dimensionality reduction and feature extraction to identify the most relevant features of the dataset. This reduction helps minimize parameters to 2*r*, which mitigates over-fitting, particularly with small datasets.

The perturbation procedure can be understood as finding electronegativity and hardness vectors *θ*_*e*_ and *θ*_*s*_ that align with the low-dimensional embedding vectors 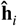 such that their inner products match the desired perturbations to *e* and *s*. Notably, any 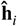 vectors in the null space of **Θ** will result in zero perturbation.

### 3.2 Fine-tuning electrostatic parameters employs efficient Zwanzig reweighting to reduce the discrepancies between espaloma-0.3.2-derived hydration free energy calculations and experimental free energies by *recycling* simulated data

With the fine-tuning mechanism in hand, we seek a loss function to minimize with respect to our data and the learnable parameters Θ of our low-rank fine-tuned correction to the foundation model (**Figure 1**). Since the objective of the investigation is to minimize the discrepancy between the experimental FreeSolv hydration free energy data and the espaloma-0.3.2 / TIP3P hydration free energy predictions, we can define a per-molecule (datapoint) discrepancy, or residual, as

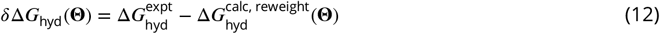

where 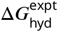 is the experimental hydration free energy, 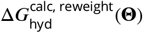 is the calculated, **Θ**-reweighted hydration free energy. The latter quantity is defined as

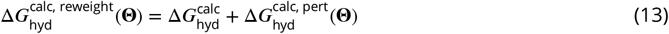

where 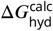 is the espaloma-0.3.2 original (i.e., unperturbed) calculated hydration free energy and its Zwanzig-reweighted perturbation [118] is given by 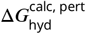 (**Θ**). From Eq. 5, the perturbation is with respect to modified small-molecule partial charges 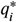, but these are a function of **Θ**, as explained in Eq. 7. More explicitly,

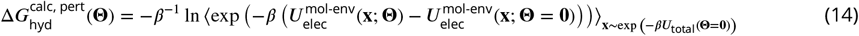

where 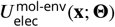 is the total molecule-environment (i.e., TIP3P solvent for hydration free energies) electrostatic potential energy (see Eq. 2) evaluated at a configuration **x** and perturbation parameters **Θ**. The first term in the exponential corresponds to the perturbed electrostatic potential whereas the second term corresponds to the unperturbed (i.e., **Θ** = **0**, or espaloma-0.3.2 from Eq. 5) electrostatic potential energy. The expectation in the Zwanzig reweighting equation is taken with respect to the molecular simulation snapshots **x** drawn from a Boltzmann distribution given by the total potential energy *U*_total_(**Θ** = **0**) in the unperturbed regime. *β* = *k*_*B*_*T*, the thermal energy of the bath. All energies and configurations are implicitly taken at the non-alchemical, fully-interacting *λ* = 1 state.

Eq. 14 implies that perturbations to the calculated (i.e., espaloma-0.3.2-derived) hydration free energies 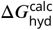 can be performed in a one-shot manner. In other words, new molecular configurations **x** need not be re-drawn from the Boltzmann distribution defined by perturbed partial charges 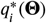 in order to compute perturbed free energies, and consequently, hydration free energy residuals *δ*Δ*G*_hyd_(**Θ**) from Eq. 12. Hence, molecular conformations from the original espaloma-0.3.2 hydration free energy calculations (see **Figure 2**) can be *recycled* to refitting molecular partial charges. Given that the most significant computational bottleneck in the entire process is the generation of uncorrelated equilibrium snapshots via molecular dynamics simulation, one-shot refitting becomes a computationally efficient fine-tuning technique since orders of magnitude fewer energy and gradient evaluations are needed for reweighting and optimization than for this initial simulation to generate the snapshots.

### 3.3 Zwanzig reweighting of free energies via molecule-solvent electrostatic interactions dramatically reduces the computational cost and memory footprint required to perform fine-tuning

Since our perturbation mechanism *exclusively* perturbs the electrostatic potential energy of the molecule-environemnt interactions (see Eq. 14), we may take advantage of the fact that the electrostatic potential energy (see Eq. 2) is linear in the molecular partial charges to dramatically reduce the amount of data which must be saved in the free energy calculation process for the fine-tuning procedure. Specifically, we need only save to disk the electrostatic potential 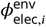 (energy per charge, in kcal (mol charge)^−1^) of the environment on each small-molecule atomic partial charge 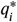 for each trajectory frame generated in the original hydration free energy calculation process rather than the full frame, itself. More explicitly,

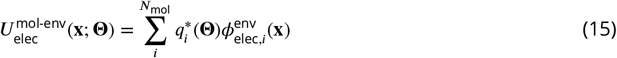

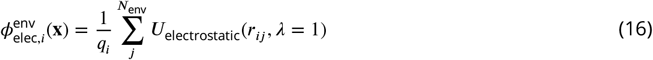

where Eq. 15 corresponds to the total molecule-environment electrostatic potential energy on all *N*_mol_ small-molecule atoms *i* and Eq. 16 corresponds to the electrostatic potential on small-molecule atom *i* from all environment atoms *j* within cutoff *r*_cut_=12Å. Only Eq. 16 for each molecular partial charge *i* needs to be saved to disk and loaded to memory rather than the entire *N*_total_-by-3 trajectory frame **x**. Eq. 15 can be computed in the fine-tuning procedure on-the-fly as an inner product with the *N*_mol_ molecular partial charges 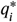 in linear time. This reduces the amount of data from the hydration free energy calculation that must be saved to disk and loaded into memory from ∼ 10^3−4^ atomic position vectors to a single 1D array of ∼ 10^1−2^ floats per frame. Consequently, all FreeSolv data necessary for Zwanzig reweighting can easily be loaded into memory for rapid fine-tuning.

## 4 Effective sample size regularization of fine-tuning via Zwanzig reweighting allows for electrostatic parameter refitting without the need to re-simulate molecular conformations by steering optimization away from uncertain free energy estimates

### 4.1 Zwanzig reweighting changes the statistical weights of the data distribution, which can cause unreliable free energy perturbation estimates when the effective sample size becomes too small

The Zwanzig reweighting term 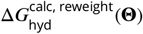 from Eq. 13 facilitates efficient fine-tuning via the recycling of pre-generated molecular conformations from the original espaloma-0.3.2 free energy calculations. However, Zwanzig reweighting has subtle limitations, creating an inherent trade-off between efficiency and both bias and precision [94]. Specifically, as the same data are recycled to minimize Eq. 12 through reweighting, the potential for erroneous optimization increases due to both high uncertainty and high bias in the estimate of 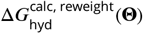. This increased error can compromise the accuracy of the refitting procedure and lead to unreliable results that do not generalize.

The cause of this higher free energy error is variance and bias accumulation, which is an inherent consequence of reweighting [51, 94]. Specifically, each time a conformation is reweighted against a new reduced potential energy surface (parameterized by **Θ**), its statistical “weight” or “importance” changes because the likelihood of observing that conformation under the new probability distribution function is modified. Reweighting adjusts the relative likelihood of observing each conformation to reflect changes in the potential energy landscape, which can either increase or decrease the effective representation of that conformation in the final statistical ensemble. As the ‘phase-space’ overlap (see **Figure 3A**, *right*), described in statistical physics parlance, or the intersection between the original and new probability distributions decreases, the variance in the free energy estimate increases [114]. The bias that accumulates upon reweighting is a direct consequence of having only a finite number of conformations to compute the free energy, and can introduce comparable order to the variance for this estimator [94]. This limited sample size reduces estimate precision and leads to bias accumulation, as it restricts the diversity of sampled conformations, making it difficult to fully capture the system’s variability. As the potential energy surface changes, the statistical weight of each conformation is altered, effectively reducing the number of independent samples. When this ‘effective’ sample size becomes too low, the Law of Large Numbers no longer holds, resulting in finite sample size effects and increased bias. Therefore, reweighting introduces uncertainty that must be carefully monitored and mitigated during fine-tuning.

**Figure 3.**
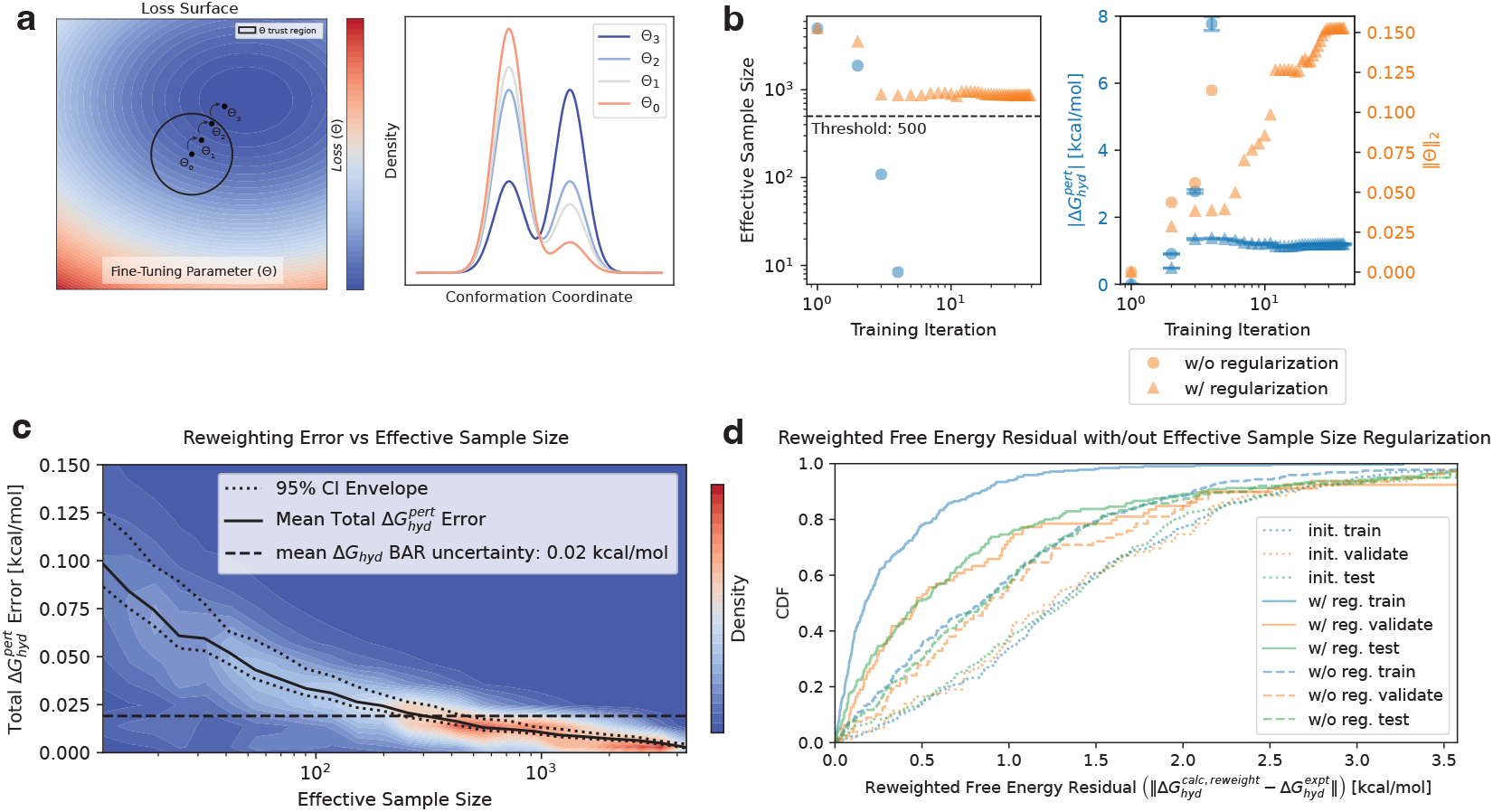
Single-shot model fine-tuning with effective sample size (ESS) regularization allows for parameter optimization via reweighting whilst maintaining a small reweighting error and avoiding sample size collapse. In order to fine-tune a model without requiring extensive re-simulation at updated parameter values, we develop a strategy to prevent ESS collapse (and therefore maintain small reweighting error) during parameter fine-tuning. **(A)** *Left*: Optimization of fine-tuning parameters **Θ** on the loss surface (cartoon, color map with contour plot) should remain inside some well-defined trust region (black circle) to avoid ESS size collapse for reweighted free energy estimates as model parameters are adjusted to fit training data. *Right*: free energy reweighting by optimizing model parameters **Θ** necessarily changes the probability density function of the conformation space of a representative small molecule, which can lead to a collapse in ESS when reweighting the original simulation data to estimate free energies at updated model parameters. **(B)** *Left*: Iterative changes in the probability density function upon reweighting cause the ESS of a molecule’s sampled conformations to fall as **Θ** approaches the boundary of the trust region. Without ESS regularization (blue circles), **Θ** may move outside the trust region and sample size collapse ensues, which would require generating new simulation data to continue reliable optimization. With ESS regularization (orange triangles), the ESS is restrained above some ESS threshold (black dotted horizontal line), enabling reliable optimization progress in directions that do not immediately lead to sample size collapse. Plot shown for FreeSolv molecule “mobley_1781152”. *Right*: Optimizing the fine-tuning parameters **Θ** to minimize training loss causes the magnitude of the free energy perturbation (blue/orange circles corresponding to with and without ESS regularization, respectively) to increase as **Θ** (*L*2 norm shown in blue/orange triangles corresponding to with and without ESS regularization) is perturbed from its initial zero matrix (at training iteration 1). **(C)** As the ESS for reweighting collapses, the combined reweighting bias and error begins to increase rapidly beyond a critical ESS threshold. We define the threshold for the acceptable mean total error using the original mean (taken over all molecules) hydration free energy calculation BAR uncertainty (0.02 kcal mol^−1^). The upper 95% CI of the total error envelope falls below the error threshold near 500 samples (10% of all frames collected for each molecule from an original sample size of 5000). **(D)** Optimization using ESS regularization affords more improvement in the residuals than optimization without regularization, as can be seen from the empirical cumulative distribution function (CDF) of absolute free energy residuals of the Optimized/Reweighted (solid lines) train/test/validate data partitions using ESS regularization as compared to that without (dashed lines). The original (unperturbed) free energy residual CDF is shown in dotted lines. Optimization without regularization is terminated when a data point falls below the ESS threshold of 500 samples as calibrated from panel **C**.

### 4.2 Effective Sample Size (ESS) serves as a heuristic to monitor the reliability of a data sample for Zwanzig reweighting of free energies

One practical approach to maintain a suitably small free energy uncertainty for reliable fine-tuning is to restrict optimization to a “trust region” in **Θ**-space (see **Figure 3A**, left), wherein the uncertainty of the reweighted free energy perturbation remains suitably small to ensure reliable results. While directly monitoring the uncertainty of 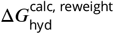 to ensure localization of the optimizer within the “trust region”, the bootstrapping required is not well-suited for automatic differentiation protocols. Instead, we use a readily-computable and differentiable proxy based on the data’s statistical weights, known as the effective sample size (ESS) [61], to monitor the reliability of our free energy estimates. For each data point (i.e., trajectory frame) **x**_*i*_ of a single molecule, the ESS as a function of fine-tuning model parameters **Θ** is given by

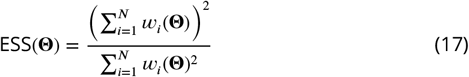

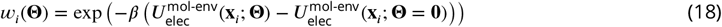

From Eq. 18, the (un-normalized) importance weight *w*_*i*_(**Θ**) of a data point is proportional to the exponential of the electrostatic potential energy difference; the same term appears in Eq. 14. This is no surprise considering that Zwanzig reweighting is a type of importance sampling (IS) [103]. From Eq. 17, the ESS of *N* samples can range from 1 to *N* (inclusively).

The effective sample size (ESS) is a commonly used heuristic for assessing the number of effectively retained samples following statistical reweighting in the Monte Carlo literature [61]. Much like the free energy uncertainty estimate, the ESS becomes unreliable in the regime of small sample sizes [54]. This unreliability arises primarily due to increased variance and the higher likelihood of over-representing a small subset of samples [8, 54]. In such cases, the effective number of statistically independent samples is often overestimated, leading to inaccurate estimates of free energy differences.

Using ESS as a proxy for reweighted 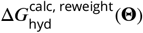 uncertainty to define a “trust region” in **Θ**-space raises a natural question: what is an appropriate ESS threshold below which a molecule’s reweighted free energy estimate can be considered unreliable? In other words, how should the ESS-derived “trust region” boundary be defined?

### 4.3 Effective sample size regularization restrains optimization to a subspace of model parameters to ensure reliable Zwanzig reweighting free energy estimates to enable one-shot fine tuning

To determine an appropriate ESS threshold to define a “trust region” boundary, a ESS-threshold calibration was performed, as shown in **Figure 3C**. In the absence of prior knowledge regarding a suitable train/test/-validation split or the optimal model rank *r*, a 50% train split was chosen with *r* = 5, corresponding roughly to the 50th percentile of cumulative variance in the PCA eigenspectrum (see **Figure 1** (insert)). The Broyden–Fletcher–Goldfarb–Shanno (BFGS) [30] algorithm minimized the ESS-unregularized loss function during optimization of the fine-tuning parameters **Θ**. For all the training molecules with final ESS values exceeding 2500 samples–a threshold chosen to prevent premature ESS collapse for too many molecules– bootstrapping was performed on the 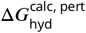 across a range of bootstrap sizes spanning an ESS range of ∼10 to 5000. The total free energy perturbation error, which includes both the Zwanzig average uncertainty 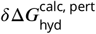 and the bias relative to the non-bootstrapped 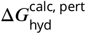 was plotted as a function of the ESS, shown in **Figure 3C**. To define the total 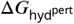 error threshold, the mean Bennett Acceptance Ratio (BAR) [7] uncertainty from the original espaloma-0.3.2 absolute hydration free energy calculation 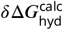 (0.02 kcal mol^−1^) was used. The corresponding ESS threshold was chosen where the upper 95% CI of the free energy perturbation uncertainty falls below this aforementioned BAR uncertainty.

In order to penalize ESSs in the loss function that fall below 500 samples, we define a *C*^∞^, piecewise flat-to-quadratic regularization function:

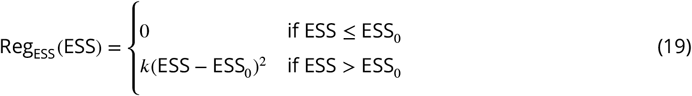

*k* and *a* are hyper-parameters that determine the stiffness of the quadratic term and the horizontal onset value of the ESS regularization. We found *k* = 100 and *a* = 750 to be sufficient to avoid the ESS-monitored early stopping. The **Θ**-dependence of the ESS is suppressed for readability, but is defined by Eq. 17.

Incorporating the ESS-regularized 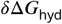 (i.e., residual between the experimental and reweighted free energy given by Eq. 12) into a differentiable loss function for optimization of fine-tuning parameters **Θ** gives the total loss function:

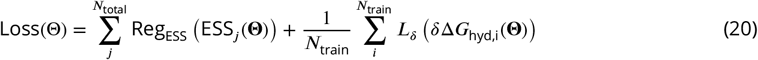

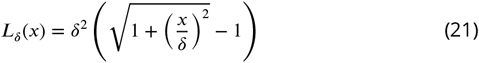

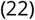

The first term in Eq. 20 represents the *sum* of the ESS penalties for all (i.e., train/test/validate) molecules *j*. ESS regularization is applied to all molecules in the interest of avoiding sample size collapse among **all** molecules. The second term penalizes free energy residuals that deviate from zero in the training set, weighted by the pseudo-Huber [46] loss function (i.e., Eq. 21). The pseudo-Huber loss is used in loss minimization since it is *C*^∞^ for smooth BFGS optimization purposes and does not over-penalize large residuals, as is the case with the *L*_2_ loss. The latter point makes the pseudo-Huber loss robust to outliers. *δ* was taken to be 1 *k*_*B*_*T*.

### 4.4 Fine-tuning with effective sample size (ESS) regularization outperforms un-regularized fine-tuning while avoiding sample size collapse

In order to compare and illustrate the behavior of ESS-regularized and un-regularized optimization, a single training molecule’s training data were extracted as a trajectory, as seen in **Figure 3B**. On the left plot, the ESS is monitored as a function of training iteration with (orange triangle) and without (blue circle). The ESS remains above the calibrated threshold of 500 (from **Figure 3C**) with regularization for nearly 50 BFGS training iteration; however, the un-regularized optimizer experiences an ESS collapse below the threshold between training iteration 2 and 3. The ESS collapse of the un-regularized optimizer on the left panel is a consequence of the relatively large perturbation of 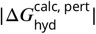 between iteration 2 and 3 (blue dot). Alternatively, the ESS-regularized 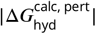 remains <2 kcal mol^−1^ for all training iterations, demonstrating that ESS regularization also curtails large free energy perturbations. Interestingly, 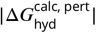 does not necessarily correlate with the ***L***_2_norm of **Θ** (orange), highlighting the nonlinear dependence of free energy perturbations on the linear electrostatic perturbation scheme. In other words, relatively large perturbations to individual electrostatic parameters does not necessarily correspond to large perturbations in the reweighted free energy.

In order to rescue loss-minimization in the unregularized control case that would otherwise experience ESS collapse as seen in **Figure 3B**, left, we imposed an early stopping when any molecule in any of the data splits achieves an ESS of <500 samples. See **Supplement** for a pseudo-code implementation of the optimization procedure.

Un-regularized refitting was compared to ESS-regularized refitting to assess whether ESS regularization actually achieves a more accurate fine-tuning model. As shown in **Figure 3D**, un-regularized (dashed), and ESS-regularized (solid) fine-tuning both improve accuracy (i.e., smaller free energy residuals) for all data splits (train/validate/test in blue, orange, and green, respectively) compared to baseline (i.e., foundation espaloma-0.3.2 free energy calculation) residuals (dotted). However, ESS-regularization consistently outperform those of the un-regularized controlled experiment in terms of accuracy, as seen from comparing the cumulative distribution function (CDF) of errors (**Figure 3D**). Interestingly, the un-regularized CDF curves tend to be consistent with each other, whereas the training ESS-regularized curve seems to separate from the validate/test curves. Based on the ESS-regularized training cluster around the zero residual, it appears that ESS-regularized optimization tends to over-fit compared to the un-regularized control. This is likely a consequence of the fact that un-regularized optimization triggers the early stopping criterion before overfitting occurs. However, ESS-regularized optimization effectively *restrains* optimization to the “trust region” without triggering early stopping. As such, the optimizer finds a local minimum that out-performs the un-regularized control. Despite over-fitting to training data in the ESS-regularized case, we find that the ESS-regularized fine-tuning model consistently out-performs the un-regularized control. This is likely because the data distribution is sufficiently consistent across all splits. This affords consistent and generalizable, albeit less dramatic, improvements in validate/test residuals correlated with improvements in training residuals. All refitting experiments were performed with a train/validate/test data split of 50/12.5/37.5% (since all refitting experiments partition validate/test data in a 25/75% split of all non-training data). Fine-tuning model rank *r* = 512 (i.e., no PCA) was chosen for ESS-regularization performance assessment as a default since no a priori knowledge about optimal rank *r* is known without calibration. This critical parameter is addressed in the following section.

## 5 Electrostatic fine-tuning demonstrates improvements in reweighted hydration free energy prediction accuracy with more training data and diminishing improvements with larger model rank

To assess the accuracy of the fine-tuned model in predicting free energies on the FreeSolv dataset, we focus on determining two critical parameters. The first is the partitioning strategy of the dataset into training, validation, and test subsets, with an emphasis on optimizing the size of the training set to determine the amount of data needed for accuracy improvements. The second parameter is the rank *r* of the fine-tuning model, which influences both dimensionality reduction and the number of parameters available for optimization. We also investigate how the model rank affects the amount of data required to achieve improvements in accuracy. All refitting experiments were performed with the ESS-regulated BFGS optimization and early-stopping criteria described in the previous section.

### 5.1 Electrostatic fine-tuning improves the accuracy of reweighted hydration free energy predictions with more training data

As shown in **Figure 4B,C**, validation and testing root mean squared errors (RMSEs) improve most significantly when increasing the training data split from 50% to 75% across all tested model ranks. While the average improvement across all data splits is positive, with a net reduction in RMSE, the largest gains occur between the 50% and 75% training data splits. Low-data regimes, involving 5% to 25% training data, also exhibit clear improvements, but these are relatively modest compared to those observed in the 75% to 90% training data ranges. Interestingly, the improvements in training data RMSE appear consistent across all data splits but only begin to align with test and validation RMSEs in the 75% to 90% range. This suggests that the optimizer tends to over-fit the training data in smaller and medium-sized data regimes.

**Figure 4.**
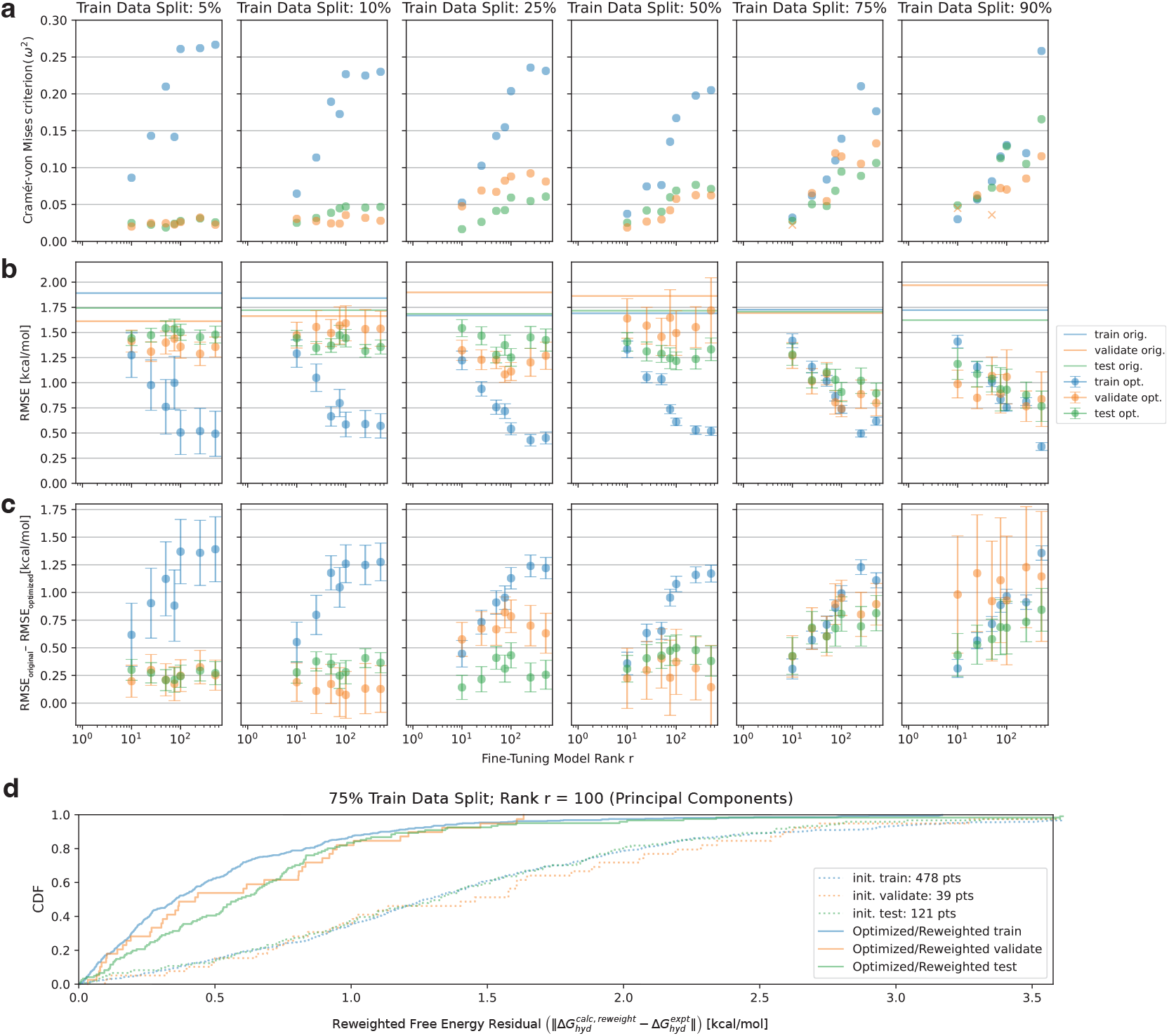
Fine-tuning experiments using a variety of training data splits and model sizes demonstrate improvements in correlation with training data and diminishing returns with fine-tuning model rank. **(A)** Cramer-von Mises criteria (ω^2^) of Optimized/Reweighted vs baseline (unoptimized) absolute free energy residual empirical cumulative distributions functions (CDFs) for a variety of training data and numbers of principal components show several trends: **(1)** test/validate residuals gradually separate with more training data; **(2)** training sets tend to separate most for all train data splits, especially for larger dimensional models; and **(3)** validate/test set separations tend to diminish with larger models beyond 100 principal components. **(B)** Reductions in RMSEs of Optimized/Reweighted free energies with respect to those of the foundation model (horizontal lines) show consistent improvement over all training splits and model ranks. **(C)** Improvements in RMSEs between Optimized/Reweighted and baseline data also show larger improvements with more training data and diminished improvements with larger model sizes. Training sets tend to overfit up to 50% training data splits. **(D)** One representative absolute free energy residual emprical cdf plot (75% training data, 100 principal components) from which the aggregate statistics in panels A,B were calculated shows significant qualitative improvement in the free energy residuals (solid lines) as compared to baselines (dotted). Model parameters Θ of this optimization experiment correspond to the largest mean improvement in validation RMSE (0.82 kcal mol^−1^) of all the experiments with >100 test data points left after data partitioning (this precludes the experiments of the rightmost column in panels **A,B,C**).

This over-fitting trend is particularly evident in **Figure 4A**, where the Cramer-von Mises statistics reveal large differences between the pre- and post-training training free energy residual CDFs (blue) as compared to those of the validation (orange) and test (green) CDFs, at least up to the 75% to 90% training data split plots. Specifically, the Cramer-von Mises [19] criterion measures the difference between pre- and post-training CDFs for each data partition (see **Figure 4D** dotted and solid CDFs, respectively, as an example) by quantifying the squared deviations across their entire range, providing a comprehensive assessment of their similarity. The explicit equation is given by

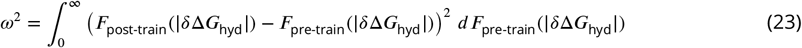

where *F* is the CDF. Importantly, nearly all the CDF comparisons reject the null hypothesis (except for two cases, marked with “x”s), with a significance level of 0.05. In this context, the null hypothesis assumes that there is no significant difference between the free energy residual CDF pre- and post-training. However, statistically significant improvements in RMSE for training partitions at a 5% split and validation partitions at 90% splits (see **Figure 4C,D** first and last columns, respectively) are difficult to discern due to the relatively small number of data points, which result in reduced statistical power and increased variability. This makes it challenging to confidently detect differences or improvements in model performance, as evident from the large error bars. This limitation is an inherent consequence of the data partitioning procedure for these extremes.

### 5.2 Electrostatic fine-tuning delivers hydration free energy accuracy improvements that diminish with model rank

In Section 3, we hypothesized that a data-driven dimensionality reduction and fine-tuning model parameterization technique would mitigate the need for large model rank. Evidence from **Figure 4B,C** supports this hypothesis to some extent, showing that while accuracy continues to improve with increasing model rank *r*, the rate of improvement diminishes significantly. Focusing on the 5%–50% data splits in **Figure 4A**, log-linear plots of model rank *r* against the Cramer-von Mises criteria reveal an increase in *ω*^2^ with *r* for all data splits that diminishes around *r*_max_ = 512. This flattening phenomenon is even more pronounced in training RMSEs, where the benefit of increasing rank diminishes noticeably. For validation and test sets in the same regime, a more prominent plateau is evident at ranks exceeding approximately 100, as the weighted square of the gap between original and optimized CDFs ceases to separate.

For RMSE improvements, the diminishing returns with larger model ranks are even more pronounced in the validation and test datasets within the 5%–50% training split regimes. By contrast, in the 75%–90% data split regimes, the Cramer-von Mises criteria (**Figure 4A**, ultimate and penultimate columns) exhibit a more log-linear trend, although over-fitting becomes evident for ranks exceeding approximately 100. A similar log-linear improvement is observed in RMSEs (**Figure 4B,C**, ultimate and penultimate columns), but again, over-fitting is apparent at higher ranks. Importantly, no statistically significant improvements are observed in test RMSEs with increased rank *r*.

The most dramatic improvement between pre- and post-fine-tuning validation RMSE for which >100 molecules (specifically 121) were present in the training partition was given by a 75% training data split with rank *r* = 100. The corresponding absolute residual CDFs are shown in **Figure 4D**. Here, over-fitting is not particularly dramatic, which is demonstrated by the relative overlapping of the train/validate/and test CDFs. We further investigate the model parameters **Θ** derived from this refitting experiment in the next section.

## 6 Optimally-fine-tuned hydration free energy predictions upon reweighting consistently recover accurate re-simulated hydration free energies

To assess the validity and reproducibility of the Zwanzig-reweighted hydration free energies predicted by the optimized fine-tuning parameters **Θ** from **Figure 4D**, we re-simulated the FreeSolv dataset’s hydration free energies using atomic partial charges 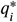 derived by the same fine-tuning parameters **Θ** for comparison. The decision to use these specific model parameters reflects a realistic experimental scenario where optimal model parameters are determined based on the best test-blind validation accuracy improvements. The decision also balanced the need to leave enough data points for testing, ensuring that prospective hydration free energy predictions could achieve tight error bounds for statistical significance.

**Figure 5A** shows the correlation plot of optimized/reweighted hydration free energy predictions against re-simulated (recalculated) hydration free energies for all molecules in the FreeSolv dataset. The RMSE of 0.06 kcal mol^−1^ highlights the precision of the reweighting procedure in recovering calculated free energies.

**Figure 5.**
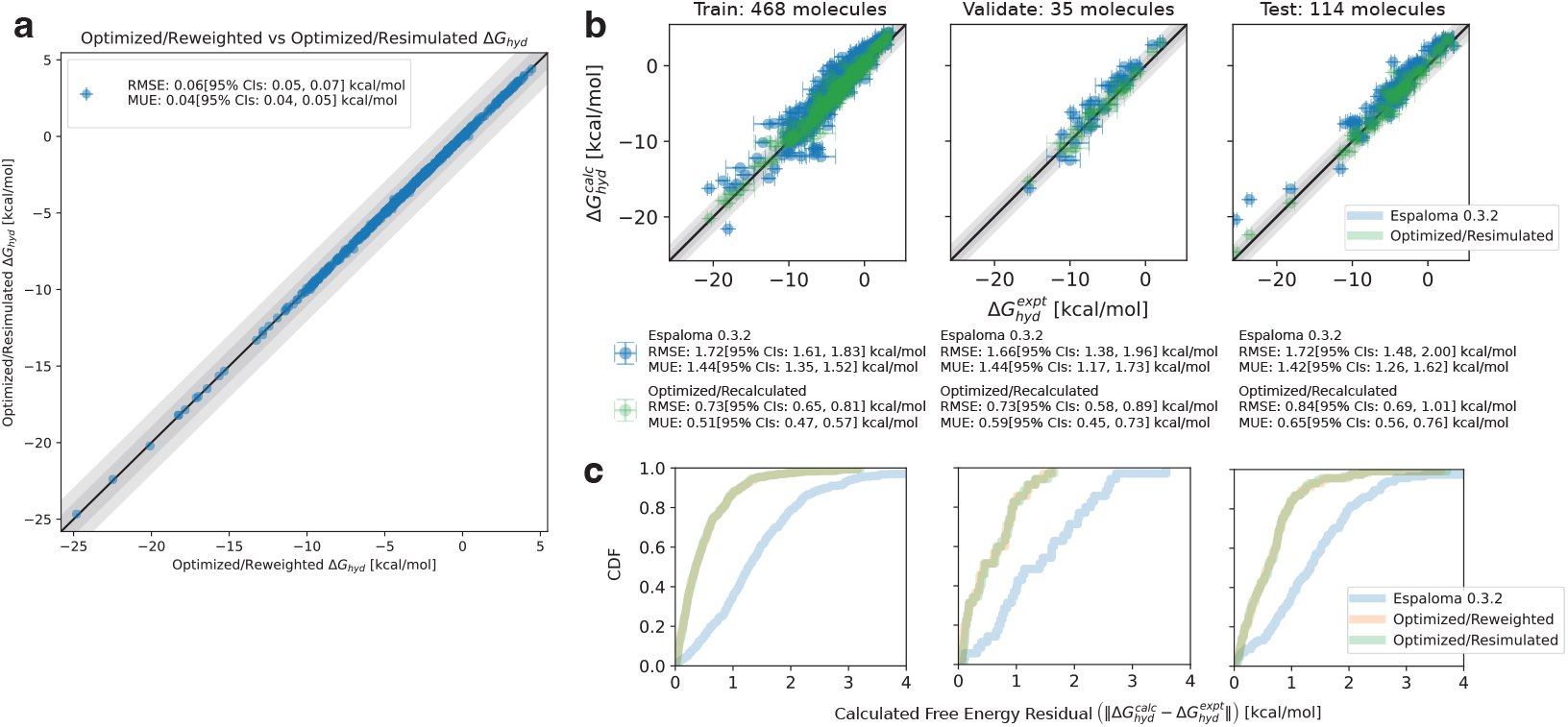
Fine-tuned Optimized/Reweighted hydration free energies are highly consistent with Optimized/Recalculated hydration free energies and demonstrate statistically significant accuracy improvements over the foundation model espaloma-0.3.2. **(A)** Correlation plot between Optimized/Reweighted free energies and Optimized/Resimulated free energies show RMSEs of 0.06 kcal mol^−1^, demonstrate agreement between Zwanzig reweighting used in fine-tuning optimization and the Bennet Acceptance Ratio (BAR) [7], which was used in the free energy recalculation experiments at optimized molecular partial charges. **(B)** Experimental vs calculated hydration free energy plots by dataset show consistent improvements in the Optimized/Resimulated calculations over the baseline foundation model espaloma-0.3.2. **(C)** Absolute hydration free energy residual CDFs for each data split and experiment show high agreement between residuals of the Optimized/Resimulated (green) and Optimized/Reweighted (orange) data (consistent with panel **A**) and reliable improvement over the baseline foundation model (green) in all data splits.

**Figure 5B** presents correlation plots comparing experimental hydration free energies to predictions from the foundation model (espaloma-0.3.2, in blue) and recalculated, optimized predictions (in green) for each data split. All splits show statistically significant improvements in RMSE and MUE compared to the foundation model. While improvements for the training set are expected, the consistent reproducibility and accuracy gains in the validation and testing sets are particularly noteworthy, underscoring their importance for reliable prospective predictions. The residual CDFs for original, reweighted, and recalculated free energy predictions across the data splits are shown in **Figure 5C** (blue, orange, and green, respectively). The high overlap between optimized/reweighted and resimulated free energy residual CDFs again demonstrates the high correlation between the two prediction methods, which is corroborated by **Figure 5A**. The improvements in the absolute residual CDFs of the optimized models over the foundation model also validates the qualitative improvements observed in **Figure 5B**.

The consistency between the optimized/reweighted and optimized/recalculated free energies, along with their significant improvement over the foundational model, suggests several key takeaways. First, the calibrated ESS threshold of 500 appears sufficiently large to mitigate the deleterious effects of finite sample sizes in the Zwanzig-calculated free energies, resulting in agreement with the recalculated BAR estimator. Additionally, ESS regularization does not seem to introduce bias in the reweighted free energy predictions when compared to the recalculated predictions. A more detailed discussion of this potential concern is provided later. Finally, improvements in free energy residuals for both weakly and strongly hydrating small molecules in the test set (as shown in the rightmost plot in **Figure 5B**) suggest that the model trained on the available data generalizes effectively to the test set, despite the sparse representation of strongly hydrating small molecules (shown in the leftmost plot in **Figure 5B**). This suggests that the fine-tuned model demonstrates robustness not only for interpolation but also for extrapolation.

## 7 Discussion

### 7.1 Fine-Tuning molecule-environment electrostatics is especially suited to memory and computational efficiency, though steric fine-tuning is feasible with some considerations

In this investigation, we focused our fine-tuning efforts exclusively on partial atomic charges, excluding other terms such as the Lennard-Jones potential. This decision stems from the well-documented polarization deficiencies in most MM force fields, which are primarily due to the electrostatics model. Within the space of 1-body terms available for fine-tuning in Class-I MM force fields, the only alternative is steric interactions. In section 3, we mentioned that fine-tuning electrostatics is highly memory-efficient due to the linear dependence of the electrostatic potential energy on atomic partial charges (see Eqs. 15, 16). Indeed, the Lennard-Jones functional form also admits a factorization with a linear dependence on *ε* and *σ* [73] under Lorentz-Berthelot combining rules, making it similarly memory-efficient. However, the repulsive term in Eq. 1 decays as 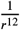. Consequently, we hypothesized that even small perturbations to the sigma or epsilon parameters in the Lennard-Jones potential would likely lead to significant decreases in the phase space overlap of our hydrated small-molecules under investigation, thereby limiting the range of optimizable parameters before ESS collapse, necessitating re-simulation.

### 7.2 Fine-Tuning is a flexible tool that is extensible to many-body terms in the MM forcefield

Fine-tuning many-body interactions, such as the bonds, angles, and torsional energetics of intra-ligand interactions, is also feasible using our dimensionality reduction procedure highlighted in blue from **Figure 1**. For 2-, 3-, and 4-body terms, we propose that an analogous data-driven dimensionality reduction and finetuning approach is achievable. Rather than concatenating one-body atom embeddings into the data matrix **H**, this approach requires using the bond, angle, and torsion embeddings generated during espaloma’s Stage 2 procedure [111]. This ensures that the many-body embeddings remain symmetric under reversal of order. At this stage, an optimizable linear transformation can be applied to compute perturbations to existing bonded parameters as was performed for the electrostatic parameters *e* and *s*. While not all of these parameters are linear in potential energy, the computational and memory demands remain low since the number of terms does not scale with the square of the number of particles, as is the case with nonbonded interactions. Instead, the number of many-body terms is approximately linear in the number of small-molecule atoms, requiring only saving the small molecule conformations from simulation trajectories for refitting.

However, it is not immediately clear whether perturbations to many-body intramolecular energetics would have a significant impact on free energy prediction accuracy. We speculate that this may be less pronounced because the foundation model already fits intramolecular interactions on high quality QM calculations of molecules in the vacuum phase [100]. Consistent with the foundation model’s adequate hydration free energy prediction accuracy (see **Figure 2**), we hypothesize that intramolecular many-body effects are already sufficiently captured by the important features of the QM data used to train the foundation model.

### 7.3 Fine-tuning is also extensible to discrete atom typing

While we deployed a fine-tuning procedure on an atom-type-free force field (i.e., espaloma-0.3.2), this approach is not strictly necessary. Most force fields include tabulated partial charges for different discrete atom types or atom groups, which can be perturbed in a similar manner to our investigation as long as the total charge constraint is respected. However, parsing partial charge assignment rules or data structures is a cumbersome task that is best avoided in our experience.

Furthermore, discrete atom-typed force fields do not allow for embedding dimensionality reduction in the same way as was pursued in this investigation. In the regime of discrete atom types, each atom embedding can be considered as a one-hot encoding. Depending on the number of atom types—and therefore the number of parameters to fit—this could present an obstacle when refitting electrostatic models to small datasets. However, if an atom-typing scheme existed where each atom is represented as a linear combination of a set of predefined atom types, our standard dimensionality reduction procedure would be applicable.

### 7.4 Linearized, perturbative fine-tuning is a parsimonious, efficient, and interpretable method of parameter refitting

In this investigation, our electrostatic fine-tuning model is linear in the learnable parameters **Θ** and perturbative to the foundation model. This approach was taken for several key reasons.

First, it is not immediately obvious that re-optimizing the entire espaloma-0.3.2 model, including the atom embeddings, would be advantageous. We postulated that the foundation model’s atom embeddings were sufficient to distinguish the relevant features of atomic chemical environments, making further reoptimization unnecessary. Furthermore, the computational effort involved in refitting the entire model (∼ 1 H100 GPU day) is rather cumbersome, as is the memory footprint of the foundation model’s QM data (approximately 20 GB [100]).

Instead, we sought to perform a minimally invasive refitting procedure, with an interpretable modification of the energetics for which error troubleshooting and diagnostic analysis would be feasible. This necessitates model interpretability. As such, we developed a dimensionality reduction procedure for parsimony and a comprehensible linear perturbation to the final outputs of the model. This ensures that the modifications are understandable and their effects traceable, allowing for easier error diagnosis and meaningful insights into model behavior. Additionally, this approach permits low computational effort (0.5–10 minutes for fine-tuning on an H100 GPU) and fast turnaround.

Furthermore, by restricting our fine-tuning to small electrostatic perturbations, we were able to achieve state-of-the-art performance on our hydration free energy predictions without the need for regularization against the foundation model’s QM data. This was possible because the small electrostatic perturbations did not deviate significantly from the original QM-derived parameters, thereby maintaining consistency without requiring additional regularization.

Finally, we sought to demonstrate that a fine-tuning apparatus of the foundation model need not be built under the constraints of the software on which the foundation model was built. While the original espaloma-0.3.2 model was built and trained with PyTorch [3], we built a lightweight JAX [12] implementation for fine-tuning, which offers several benefits. JAX’s NumPy [42]-like API, functional programming style, and efficient support for automatic differentiation make it particularly well-suited for rapid experimentation and optimization.

Altogether, we show that fine-tuning of foundation models can be lightweight, parsimonious, computationally efficient, and interpretable.

### 7.5 Effective sample size diagnostics for sample quality in Zwanzig-based reweighting is useful but has several drawbacks that often require careful consideration

In the previous section, we detailed the ESS-regularized fine-tuning procedure and the improvements in predictive accuracy that it affords despite the over-fitting tendencies it has. As mentioned previously, ESS is used as a convenient proxy for reweighted free energy prediction uncertainty caused by reduced phase space overlap and finite sample size bias. It provides a measure of how well the sampled conformational space represents the target distribution, making it particularly useful in indicating the reliability of free energy estimates. High ESS values suggest that the sample has sufficient effective independent information, reducing the impact of biases due to correlations or insufficient sampling, thus providing a good gauge for the robustness of reweighted predictions. However, while ESS is a particularly convenient proxy that proved to be effective in regularization to restrain the parameter optimization to an appropriate trust region, there are several drawbacks of the ESS that are worth mentioning. High ESS can occasionally mask structural problems in the target statistical ensemble under investigation. These problems include insufficient coverage of important regions of conformational space that become available upon reweighting to a new probability distribution [61, 103]. High ESS may give the false impression that the sample is comprehensive, even when crucial areas of the conformational space are not adequately represented. ESS is also sensitive to outliers in the weight distribution, which can lead to over/underestimation of the true ESS in a finite sample size regime [1]. Furthermore, ESS can experience significant bias in small sample sizes, which are under-represented if samples are correlated (i.e., not i.i.d). Consequently, applications of ESS in the Monte Carlo literature occasionally use Kullback-Leibler (KL) [56] divergence or the Geweke [9] diagnostic when computationally tractable [103]. When this is not feasible, resampling and Leave-One-Out Cross-Validation (LOO-CV) [113] are used to corroborate the ESS. Resampling methods help evaluate the stability of ESS by repeatedly drawing subsets of data, while LOO-CV assesses the robustness of the model by systematically leaving out each data point, ensuring that the ESS estimate is consistent across different sub-samples of the data. The alternative methods, however, were overlooked in the interest of computational feasibility and the need to efficiently perform autograd for loss minimization.

One other subtle pitfall of ESS regularization in the refitting procedure involves sample pruning. Specifically, we considered the possibility that minimizing a loss function with ESS regularization might, for a given molecule, select a sub-ensemble of molecular conformations that consistently have high weights and small free energy residuals. However, these conformations may lack sufficient coverage of the underlying probability distribution function, leading to an incomplete representation of the ensemble. This lack of coverage can be problematic for the optimization process, as it may result in biased free energy estimates and reduced generalizability of the model. We anticipate this becomes more of a possibility as the flexibility of the underlying fine-tuning model increases or the ESS threshold drops. Increased model flexibility can lead to overfitting specific high-weight conformations, while a lower ESS threshold allows the model to rely on fewer effective samples, both of which can exacerbate the issue of insufficient coverage of the conformational space.

In this study, the aforementioned pitfalls of ESS and ESS-regularized refitting were avoided to recover accurate reweighted free energy estimates compared to re-simulation and experimental predictions (see **Figure 5**). We suspect this is a consequence of having used a rather conservative ESS threshold for regularization (500 samples), which corresponded to a mean total reweighted free energy uncertainty of 0.02 kcal mol^−1^. It is certainly possible that using a more aggressive reweighting strategy by reducing the ESS threshold further might afford even better hydration free energy predictions at the cost of slightly higher uncertainty. We leave this to a future investigation.

### 7.6 Effective sample size regularization across all data splits is necessary to avoid sample size collapse in validation and test sets and suits prospective molecular property prediction

In Eq. 20, ESS regularization is applied to **all** train, validation, and test data sets. It was mentioned that this precaution was taken to avoid spurious free energy residual minimization upon sample size collapse in the training data and to retain all possible test set predictions. We argue that using ESS regularization across all data splits is an appropriate methodology that is consistent with and can be incorporated into the computational molecular property prediction pipeline.

In computational molecular property prediction pipelines, one might imagine first collecting experimental data, then using an existing foundation force field to perform simulations and subsequent predictions (e.g., hydration free energy predictions). Then molecules with the best properties (e.g., hydration free energies, binding affinities, ADME properties) are selected for synthesis and assay. However, since synthesis and assay times are not consistent for all molecules, data streaming ensues. Asynchronously, assay data may be integrated into the pool of training data, simulated at the most recently fine-tuned force field model, and fine-tuned on-the-fly for subsequent property predictions on newly recommended molecules. Importantly, ESS regularization avoids loss of property prediction accuracy on test molecules, meaning there is no need to introduce a re-simulation bottleneck on prospective molecules. In fact, any molecules for which simulation data exist at any force field can be reweighted to the most recent fine-tuned model to inform downstream synthesis and validation through assaying. Hence, ESS regularization on the test set is advisable to avoid deterioration of prospective molecular property predictions and increase the efficiency of molecular property optimization.

## 8 Conclusion

In this investigation, we demonstrate that fine-tuning the espaloma-0.3.2 foundation charge model to refit experimental hydration free energies using a low-rank, Zwanzig-reweighting scheme achieves state-of-the-art accuracy among MM force fields. This approach utilizes data-driven dimensionality reduction of atomic embeddings and linear perturbation of foundation-model electrostatic parameters, resulting in a parsimonious and interpretable refitting procedure. Perturbed electrostatic parameters are used to recompute small-molecule atomic partial charges via the QEq scheme, maintaining the total molecular charge while perturbing the molecule-solvent electrostatic potential energy. By focusing solely on perturbing the molecule-solvent electrostatic potential energy, we present a data- and compute-efficient methodology that effectively minimizes the residual between experimental free energies and those computed from the foundation model.

Zwanzig reweighting of foundation-model absolute hydration free energy-derived molecular electrostatic potentials, combined with effective sample size (ESS) regularization, provides an efficient means to optimize fine-tuning electrostatic parameters without the need for re-simulating molecular conformations. Specifically, ESS regularization restrains the fine-tuning model parameter space to a trust region of reliably reweighted free energies, mitigating the risk of catastrophic sample size collapse. By preventing erroneous minimization into unreliable regions of parameter space, ESS regularization preserves the model’s predictive capacity and avoids premature early stopping of the optimization procedure.

Our results consistently show that fine-tuning improves free energy accuracy, with some noteworthy trends. The accuracy improvement is most dramatic in the 75%-90% training split regime, where the model benefits from sufficient data for optimization. However, accuracy improvements diminish with larger finetuning model ranks, particularly in the small data regime, suggesting that increased model complexity does not yield significant accuracy gains beyond a model rank of ∼ 100.

Predicted hydration free energies via optimally fine-tuned Zwanzig reweighting show a high correlation with the re-simulated absolute hydration at the same optimized model parameters, **Θ**. One-shot Zwanzig reweighting proves to be a robust and accurate methodology that is statistically indistinguishable from the more computationally intensive method of re-simulating free energies at optimized parameters. This demonstrates that Zwanzig reweighting provides a highly efficient alternative to traditional re-simulation approaches while maintaining accuracy.

Such a lightweight fine-tuning procedure is likely extensible to other reliable experimental datasets that can be accompanied by calculated predictions, such as protein-ligand binding free energies for preclinical drug discovery. We also anticipate that the one-shot fine-tuning strategy will extend to a broader class of molecular potentials, including polarizable models and neural network potentials. By leveraging this approach, potential parameters can be refined efficiently, improving the accuracy of predictions in drug and material discovery applications, where precise modeling of molecular interactions is crucial for identifying promising candidates and chemical matter with novel properties.

## 9 Disclaimer

The content is solely the responsibility of the authors and does not necessarily represent the official views of the National Institutes of Health.

### 9.1 Funding

JDC acknowledges support from NIH grant P30 CA008748, NIH grant R01 GM132386, NIH grant R35 GM152017, and the Sloan Kettering Institute. DAR acknowledges support from Relay Therapeutics and the Sloan Kettering Institute.

### 9.2 Disclosures

JDC is a current member of the Scientific Advisory Board of OpenEye Scientific Software. JDC has equity in and serves as the Chief Executive Officer of Achira, Inc., which is engaged in the creation of open foundation simulation models for drug discovery. The Chodera laboratory receives or has received funding from multiple sources, including the National Institutes of Health, the National Science Foundation, the Parker Institute for Cancer Immunotherapy, Relay Therapeutics, Entasis Therapeutics, Silicon Therapeutics, EMD Serono (Merck KGaA), AstraZeneca, Vir Biotechnology, Bayer, XtalPi, In3terline Therapeutics, the Molecular Sciences Software Institute, the Starr Cancer Consortium, the Open Force Field Consortium, Cycle for Survival, a Louis V. Gerstner Young Investigator Award, and the Sloan Kettering Institute. A complete funding history for the Chodera lab can be found at http://choderalab.org/funding.

## 10 Author Contributions

Conceptualization: DAR, JDC, JF;

Methodology: DAR, JDC, JF;

Software: DAR, JF;

Investigation: DAR;

Writing–Original Draft: DAR, JDC, JF;

Funding Acquisition: JDC, JF;

Resources: JDC;

Supervision: JDC, JF;

## 11 Acknowledgments

The authors are grateful for the support of the timemachine team, including Yutong Zhao (ORCID: 0009-0007-1527-6287), Joseph Kaus, Matthew Wittmann, and Forrest York. The authors thank Marcus Wieder for his insights into refitting potentials to experimental data and effective sample size penalization. The authors also thank Demetri Moustakas (ORCID: 0009-0000-1302-4042) for his patient mentorship and guidance during early phases of the project that led to this publication. We are grateful to OpenEye, Cadence Molecular Sciences for providing a free academic research license to the OpenEye Toolkit for this work. The authors are grateful to W. Patrick Walters and Mark Murcko for their mentorship and guidance during the course of this project.

## A Supporting Information

### A.1 Code Availability

The Python, C++, and CUDA code used to perform hydration free energy experiments and fine-tuning in this paper is distributed open source under Apache License, Version 2.0 at https://github.com/dominicrufa/timemachine as a fork from https://github.com/proteneer/timemachine. Core dependencies include PyTorch 2.0.0 [77], Deep Graph Library 0.6.0 [109], the Open Force Field Toolkit 0.11.2 [67], JAX 0.4.30 [12], and timemachine [117].

The submission scripts and notebooks associated with data generation, analysis, and visualization are found at https://github.com/dominicrufa/timemachine/tree/master/data

**Figure S 6.**
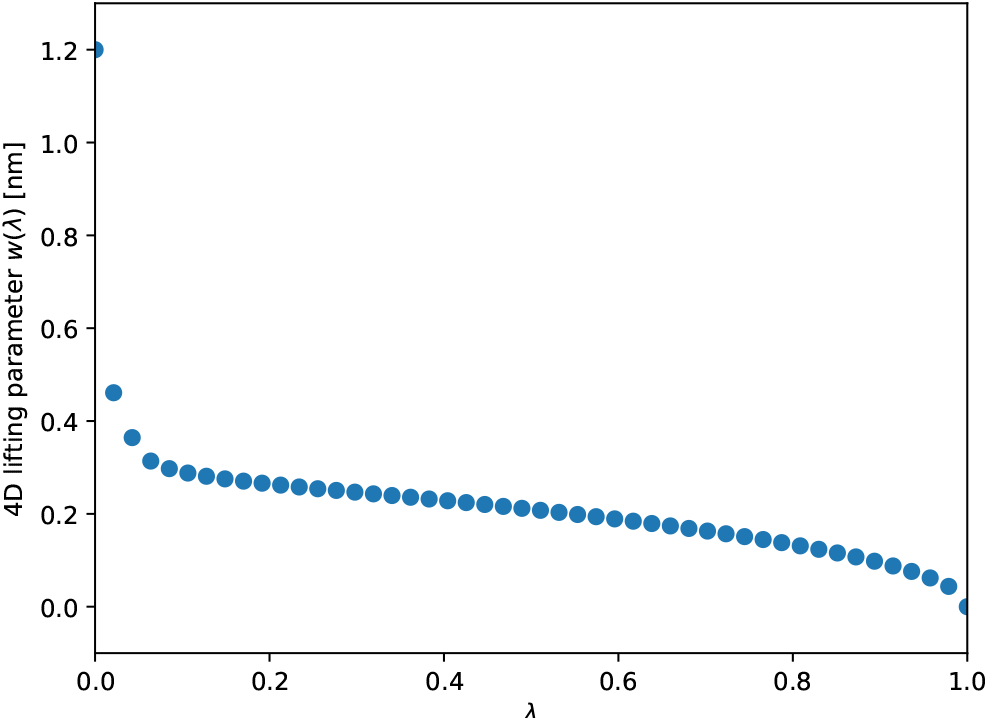
4D coupling protocol *w*(*λ*) as a function of *λ*. *λ* = 0 corresponds to the decoupled state wherein the small molecule is non-interacting with TIP3P solvent as the 4-dimensional decoupling dimension is equal to *r*_cut_. The 4-dimensional distance shrinks as *λ* progresses to 1, where no interactions are lifted.

### A.2 Detailed Methods

#### MM force field implementation

Bond, angle, and proper torsion parameters for FreeSolv small molecules were generated from the espaloma-0.3.2 foundation model and passed to the appropriate timemachine [117] energy/force evaluation functions. Openff-2.1.0 (“Sage”) [27] improper torsions were used to replace espaloma-0.3.2 impropers due to conflicting conventions in atom ordering.

#### FreeSolv hydration free energy calculation protocol

To compute hydration free energies for the FreeSolv dataset [67], we used a modified version of the protocol described in [70].

Neutral molecules were solvated with TIP3P water [50] in cubical boxes of width 4nm. Hydrogen Mass Repartitioning [44] was implemented between all heavy atoms and hydrogens by subtracting 2 amus from the former and adding to the latter.

A palindromic BAOAB Langevin integrator [57–59] was used with a friction coefficient of 1 ps^−1^ and a timestep of 2.5 fs. A Monte Carlo Barostat at a temperature of 300 K and a pressure of 1.013 bar was used which alternated with MD steps.

Lambda independent dynamics were used to evaluate free energies between *λ* = 0 and 1 using Bennett Acceptance Ratio (BAR) differences between adjacent windows. Each window ran 5000 frames (at 400 timesteps per frame saving frequency) preceded by 10,000 equilibration timesteps with 48 *λ* windows. The lifting term, *w*(*λ*), in Eq. 4 is given by **Figure 6**, which is a pre-calibrated coupling schedule

### A.3 Fine-Tuning by Zwanzig reweighting

Algorithm 1 describes the ESS monitoring procedure of BFGS optimization associated with the grid of ESS-regularized, Zwanzig-reweighted fine-tuning refitting experiments in **Figure 7**. gtol is the default value set by `scipy.optimize.minimize` with `method = ‘BFGS’`.

**Figure S 7.**
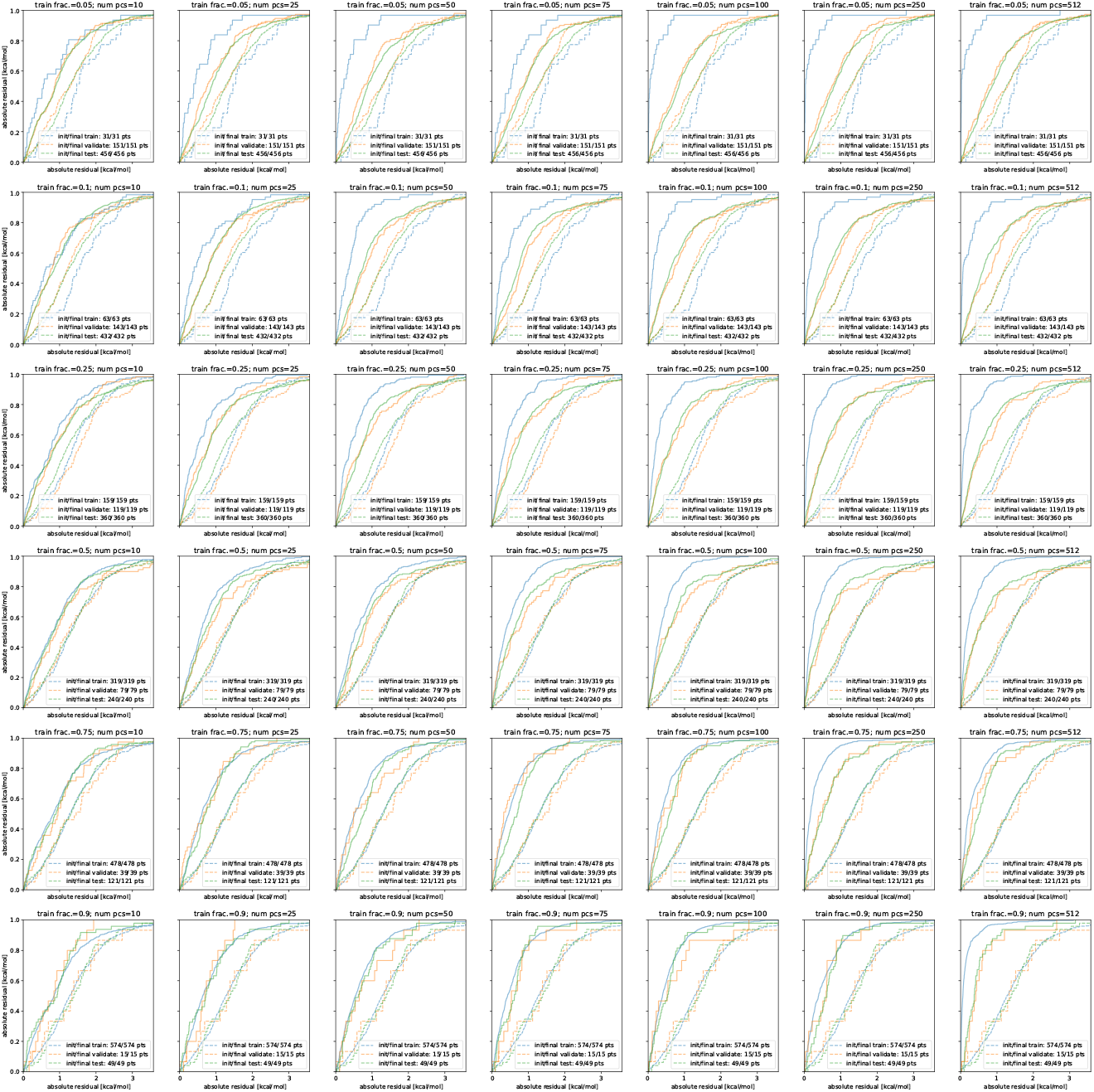
Full empirical CDFs of each ESS-regularized, Zwanzig-reweighted fine-tuning experiment. The aggregated RMSD and Cramer-von Mises statistics for each fine-tuning experiment are computed from the CDFs shown above. Training fractions and the number of PCs are depicted as graph titles above each plot. The number of data points pre- and post-optimization for each data partition that satisfy the ESS criterion are shown. Importantly, ESS regularization retains **all** data points in every partition for all experiments. The rightmost column (i.e., 512 principal components) did not include the change-of-base procedure from Eq. 6 since the dimension of each embedding is already of dimension 512 (i.e., *r*_max_)

#### Algorithm 1

BFGS optimization with effective sample size (ESS) monitoring

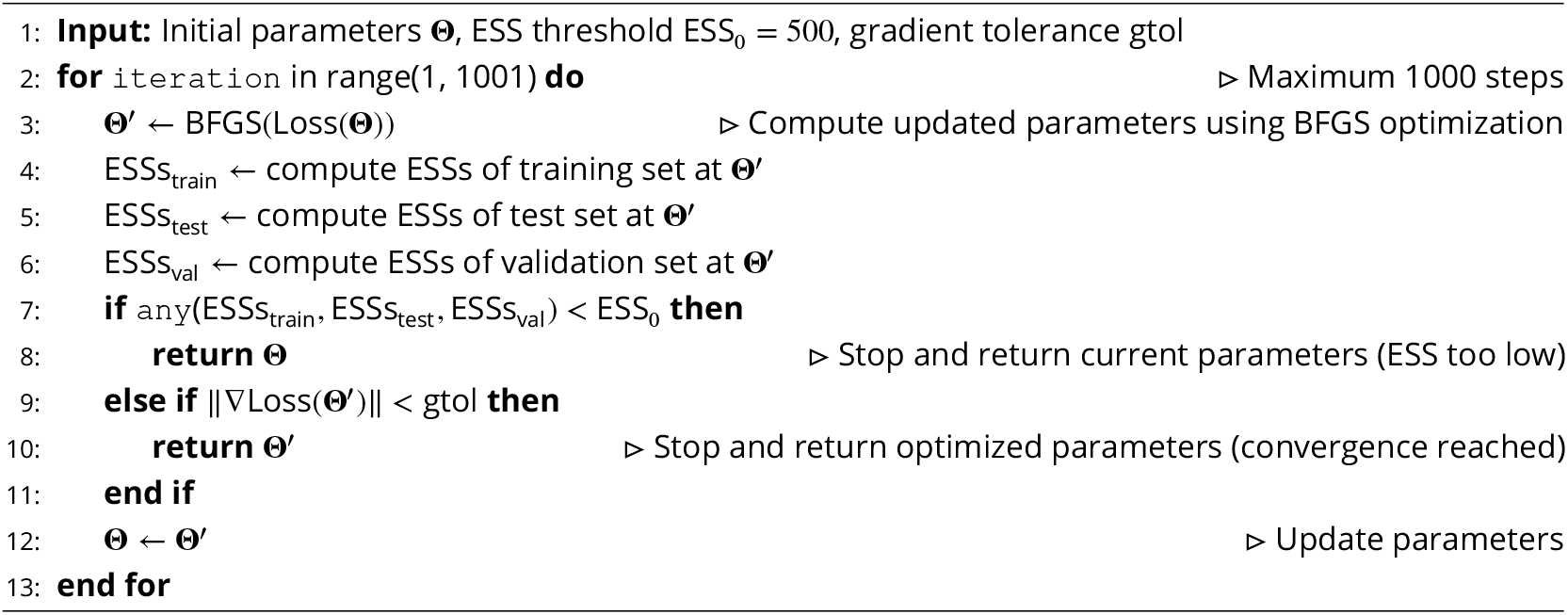

## References

[1] Aguinis, H., Gottfredson, R. K., and Joo, H. (2013). Best-practice recommendations for defining, identifying, and handling outliers. Organizational research methods, 16(2):270–301.

[2] Albanese, S. K., Chodera, J. D., Volkamer, A., Keng, S., Abel, R., and Wang, L. (2020). Is structure-based drug design ready for selectivity optimization? Journal of chemical information and modeling, 60(12):6211–6227.

[3] Ansel, J., Yang, E., He, H., Gimelshein, N., Jain, A., Voznesensky, M., Bao, B., Bell, P., Berard, D., Burovski, E., Chauhan, G., Chourdia, A., Constable, W., Desmaison, A., DeVito, Z., Ellison, E., Feng, W., Gong, J., Gschwind, M., Hirsh, B., Huang, S., Kalambarkar, K., Kirsch, L., Lazos, M., Lezcano, M., Liang, Y., Liang, J., Lu, Y., Luk, C., Maher, B., Pan, Y., Puhrsch, C., Reso, M., Saroúm, M., Siraichi, M. Y., Suk, H., Suo, M., Tillet, P., Wang, E., Wang, X., Wen, W., Zhang, S., Zhao, X., Zhou, K., Zou, R., Mathews, A., Chanan, G., Wu, P., and Chintala, S. (2024). Pytorch 2: Faster machine learning through dynamic python bytecode transformation and graph compilation. In Proceedings of the 29th ACM International Conference on Architectural Support for Programming Languages and Operating Systems, Volume 2 (ASPLOS ‘24). ACM.

[4] Anstine, D., Zubatyuk, R., and Isayev, O. (2024). Aimnet2: a neural network potential to meet your neutral, charged, organic, and elemental-organic needs.

[5] Bartlett, R. J. and Musiał, M. (2007). Coupled-cluster theory in quantum chemistry. Reviews of Modern Physics, 79(1):291–352.

[6] Batzner, S., Musaelian, A., Sun, L., Geiger, M., Mailoa, J. P., Kornbluth, M., Molinari, N., Smidt, T. E., and Kozinsky, B. (2022). E (3)-equivariant graph neural networks for data-efficient and accurate interatomic potentials. Nature communications, 13(1):2453.

[7] Bennett, C. H. (1976). Efficient estimation of free energy differences from monte carlo data. Journal of Computational Physics, 22(2):245–268.

[8] Berger, J., Bayarri, M., and Pericchi, L. (2014). The effective sample size. Econometric Reviews, 33(1-4):197–217.

[9] Bernardo, J., Berger, J., Dawid, A., and Smith, A. (1992). John geweke. In Bayesian Statistics 4: Proceedings of the Fourth Valencia International Meeting: Dedicated to the Memory of Morris H. DeGroot, 1931-1989: April 15-20, 1991, volume 4, page 169. Clarendon Press.

[10] Boothroyd, S., Behara, P. K., Madin, O. C., Hahn, D. F., Jang, H., Gapsys, V., Wagner, J. R., Horton, J. T., Dotson, D. L., Thompson, M. W., et al. (2023). Development and benchmarking of open force field 2.0. 0: the sage small molecule force field. Journal of chemical theory and computation, 19(11):3251–3275.

[11] Boothroyd, S., Madin, O. C., Mobley, D. L., Wang, L.-P., Chodera, J. D., and Shirts, M. R. (2022). Improving force field accuracy by training against condensed-phase mixture properties. Journal of chemical theory and computation, 18(6):3577–3592.

[12] Bradbury, J., Frostig, R., Hawkins, P., Johnson, M. J., Leary, C., Maclaurin, D., Necula, G., Paszke, A., VanderPlas, J., Wanderman-Milne, S., and Zhang, Q. (2018). JAX: composable transformations of Python+NumPy programs.

[13] Case, D. A., Darden, T. A., Cheatham, T. E., Simmerling, C. L., Wang, J., Duke, R. E., Luo, R., Crowley, M., Walker, R. C., Zhang, W., et al. (2008). Amber 10.

[14] Cavasotto, C. N. (2020). Binding free energy calculation using quantum mechanics aimed for drug lead optimization. Quantum mechanics in drug discovery, pages 257–268.

[15] Chatterjee, P., Sengul, M. Y., Kumar, A., and MacKerell Jr, A. D. (2022). Harnessing deep learning for optimization of lennard-jones parameters for the polarizable classical drude oscillator force field. Journal of chemical theory and computation, 18(4):2388–2407.

[16] Chen, G., Jaffrelot Inizan, T., Plé, T., Lagardere, L., Piquemal, J.-P., and Maday, Y. (2024). Advancing force fields parameterization: A directed graph attention networks approach. Journal of Chemical Theory and Computation, 20(13):5558– 5569.

[17] Chodera, J. D., Mobley, D. L., Shirts, M. R., Dixon, R. W., Branson, K., and Pande, V. S. (2011). Alchemical free energy methods for drug discovery: Progress and challenges. Current Opinion in Structural Biology, 21(2):150–160.

[18] Cournia, Z., Allen, B., and Sherman, W. (2017). Relative binding free energy calculations in drug discovery: recent advances and practical considerations. Journal of chemical information and modeling, 57(12):2911–2937.

[19] Cramér, H. (1928). On the composition of elementary errors: First paper: Mathematical deductions. Scandinavian Actuarial Journal, 1928(1):13–74.

[20] Dauber-Osguthorpe, P. and Hagler, A. T. (2019). Biomolecular force fields: where have we been, where are we now, where do we need to go and how do we get there? Journal of computer-aided molecular design, 33(2):133–203.

[21] Dodda, L. S., Vilseck, J. Z., Tirado-Rives, J., and Jorgensen, W. L. (2017). 1.14* cm1a-lbcc: localized bond-charge corrected cm1a charges for condensed-phase simulations. The Journal of Physical Chemistry B, 121(15):3864–3870.

[22] Duarte Ramos Matos, G. (2018). Free energy calculations in action: Theory, applications and challenges of solvation free energies.

[23] Duarte Ramos Matos, G., Kyu, D. Y., Loeffler, H. H., Chodera, J. D., Shirts, M. R., and Mobley, D. L. (2017). Approaches for calculating solvation free energies and enthalpies demonstrated with an update of the freesolv database. Journal of Chemical & Engineering Data, 62(5):1559–1569.

[24] Eastman, P., Behara, P. K., Dotson, D. L., Galvelis, R., Herr, J. E., Horton, J. T., Mao, Y., Chodera, J. D., Pritchard, B. P., Wang, Y., et al. (2023a). Spice, a dataset of drug-like molecules and peptides for training machine learning potentials. Scientific Data, 10(1):11.

[25] Eastman, P., Galvelis, R., Peláez, R. P., Abreu, C. R., Farr, S. E., Gallicchio, E., Gorenko, A., Henry, M. M., Hu, F., Huang, J., et al. (2023b). Openmm 8: molecular dynamics simulation with machine learning potentials. The Journal of Physical Chemistry B, 128(1):109–116.

[26] Eastman, P., Pritchard, B. P., Chodera, J. D., and Markland, T. E. (2024). Nutmeg and spice: models and data for biomolecular machine learning. Journal of Chemical Theory and Computation, 20(19):8583–8593.

[27] Eastwood, J. B., Behara, P., and contributors (2023). openff-2.1.0 (sage). Posted on 17 Jul 2023.

[28] Ehlert, S., Stahn, M., Spicher, S., and Grimme, S. (2021). Robust and efficient implicit solvation model for fast semiempirical methods. Journal of Chemical Theory and Computation, 17(7):4250–4261.

[29] Fass, J., York, F., Wittmann, M., Kaus, J., and Zhao, Y. (2023). Local resampling trick for focused molecular dynamics. Journal of Chemical Theory and Computation, 19(18):6139–6150.

[30] Fletcher, R. (2000). Practical methods of optimization. John Wiley & Sons.

[31] Fuchs, P., Thaler, S., Röcken, S., and Zavadlav, J. (2024). chemtrain: Learning deep potential models via automatic differentiation and statistical physics. arXiv preprint 2408.15852.

[32] Galvelis, R., Varela-Rial, A., Doerr, S., Fino, R., Eastman, P., Markland, T. E., Chodera, J. D., and De Fabritiis, G. (2023). Nnp/mm: accelerating molecular dynamics simulations with machine learning potentials and molecular mechanics. Journal of chemical information and modeling, 63(18):5701–5708.

[33] Geerlings, P., De Proft, F., and Langenaeker, W. (2003). Conceptual density functional theory. Chemical reviews, 103(5):1793–1874.

[34] Gilson, M. K., Gilson, H. S., and Potter, M. J. (2003). Fast assignment of accurate partial atomic charges: an electronegativity equalization method that accounts for alternate resonance forms. Journal of chemical information and computer sciences, 43(6):1982–1997.

[35] Gräter, F., Schwarzl, S. M., Dejaegere, A., Fischer, S., and Smith, J. C. (2005). Protein/ligand binding free energies calculated with quantum mechanics/molecular mechanics. The Journal of Physical Chemistry B, 109(20):10474–10483.

[36] Greener, J. G. (2024). Differentiable simulation to develop molecular dynamics force fields for disordered proteins. Chemical Science, 15(13):4897–4909.

[37] Hafner, J. (2008). Ab-initio simulations of materials using vasp: Density-functional theory and beyond. Journal of computational chemistry, 29(13):2044–2078.

[38] Hagler, A. T. (2018). Biomolecular force fields: Where have we been, where are we now, where do we need to go and how do we get there? Journal of Computer-Aided Molecular Design, 32(8):935–939.

[39] Hagler, A. T. (2019). Force field development phase ii: Relaxation of physics-based criteria… or inclusion of more rigorous physics into the representation of molecular energetics. Journal of computer-aided molecular design, 33(2):205– 264.

[40] Hahn, D. F., Bayly, C. I., Boby, M. L., Macdonald, H. E. B., Chodera, J. D., Gapsys, V., Mey, A. S., Mobley, D. L., Benito, L. P., Schindler, C. E., et al. (2022). Best practices for constructing, preparing, and evaluating protein-ligand binding affinity benchmarks [article v1. 0]. Living journal of computational molecular science, 4(1).

[41] Harder, E., Damm, W., Maple, J., Wu, C., Reboul, M., Xiang, J. Y., Wang, L., Lupyan, D., Dahlgren, M. K., Knight, J. L., et al. (2016). Opls3: a force field providing broad coverage of drug-like small molecules and proteins. Journal of chemical theory and computation, 12(1):281–296.

[42] Harris, C. R., Millman, K. J., van der Walt, S. J., Gommers, R., Virtanen, P., Cournapeau, D., Wieser, E., Taylor, J., Berg, S., Smith, N. J., Kern, R., Picus, M., Hoyer, S., van Kerkwijk, M. H., Brett, M., Haldane, A., del Río, J. F., Wiebe, M., Peterson, P., Gérard-Marchant, P., Sheppard, K., Reddy, T., Weckesser, W., Abbasi, H., Gohlke, C., and Oliphant, T. E. (2020). Array programming with NumPy. Nature, 585:357–362.

[43] Hauser, K., Negron, C., Albanese, S. K., Ray, S., Steinbrecher, T., Abel, R., Chodera, J. D., and Wang, L. (2018). Predicting resistance of clinical abl mutations to targeted kinase inhibitors using alchemical free-energy calculations. Communications biology, 1(1):70.

[44] Hopkins, C. W., Le Grand, S., Walker, R. C., and Roitberg, A. E. (2015). Long-time-step molecular dynamics through hydrogen mass repartitioning. Journal of chemical theory and computation, 11(4):1864–1874.

[45] Hu, H., Lu, Z., and Yang, W. (2007). Fitting molecular electrostatic potentials from quantum mechanical calculations. Journal of chemical theory and computation, 3(3):1004–1013.

[46] Huber, P. J. (1992). Robust estimation of a location parameter. In Breakthroughs in statistics: Methodology and distribution, pages 492–518. Springer.

[47] Jakalian, A., Bush, B. L., Jack, D. B., and Bayly, C. I. (2000). Fast, efficient generation of high-quality atomic charges. am1-bcc model: I. method. Journal of computational chemistry, 21(2):132–146.

[48] Jakalian, A., Jack, D. B., and Bayly, C. I. (2002). Fast, efficient generation of high-quality atomic charges. am1-bcc model: Ii. parameterization and validation. Journal of computational chemistry, 23(16):1623–1641.

[49] Jambeck, J. P. and Lyubartsev, A. P. (2013). Another piece of the membrane puzzle: extending slipids further. Journal of chemical theory and computation, 9(1):774–784.

[50] Jorgensen, W. L., Chandrasekhar, J., Madura, J. D., Impey, R. W., and Klein, M. L. (1983). Comparison of simple potential functions for simulating liquid water. The Journal of chemical physics, 79(2):926–935.

[51] Jorgensen, W. L. and Thomas, L. L. (2008). Perspective on free-energy perturbation calculations for chemical equilibria. Journal of chemical theory and computation, 4(6):869–876.

[52] Kaminski, G. A., Friesner, R. A., Tirado-Rives, J., and Jorgensen, W. L. (2001). Evaluation and reparametrization of the opls-aa force field for proteins via comparison with accurate quantum chemical calculations on peptides. The Journal of Physical Chemistry B, 105(28):6474–6487.

[53] Karwounopoulos, J., Kaupang, Å., Wieder, M., and Boresch, S. (2023). Calculations of absolute solvation free energies with transformato application to the freesolv database using the cgenff force field. Journal of Chemical Theory and Computation, 19(17):5988–5998.

[54] Kolaczyk, E. D. and Krivitsky, P. N. (2015). On the question of effective sample size in network modeling: An asymptotic inquiry. Statistical science: a review journal of the Institute of Mathematical Statistics, 30(2):184.

[55] Kovács, D. P., Moore, J. H., Browning, N. J., Batatia, I., Horton, J. T., Kapil, V., Witt, W. C., Magdău, I.-B., Cole, D. J., and Csányi, G. (2023). Mace-off23: Transferable machine learning force fields for organic molecules. arXiv preprint 2312.15211.

[56] Kullback, S. and Leibler, R. A. (1951). On information and sufficiency. The annals of mathematical statistics, 22(1):79– 86.

[57] Leimkuhler, B. and Matthews, C. (2013a). Rational construction of stochastic numerical methods for molecular sampling. Applied Mathematics Research eXpress, 2013(1):34–56.

[58] Leimkuhler, B. and Matthews, C. (2013b). Robust and efficient configurational molecular sampling via langevin dynamics. The Journal of chemical physics, 138(17):05B601_1.

[59] Leimkuhler, B. and Matthews, C. (2016). effcient molecular dynamics using geodesic integration and solvent–solute splitting. Proceedings of the Royal Society A: Mathematical, Physical and Engineering Sciences, 472(2189):20160138.

[60] Lennard-Jones, J. E. (1931). Cohesion. Proceedings of the Physical Society, 43(5):461.

[61] Lenth, R. V. (2001). Some practical guidelines for effective sample size determination. The american statistician, 55(3):187–193.

[62] Lu, X., Fang, D., Ito, S., Okamoto, Y., Ovchinnikov, V., and Cui, Q. (2016). Qm/mm free energy simulations: recent progress and challenges. Molecular simulation, 42(13):1056–1078.

[63] Lundborg, M., Wennberg, C., Lidmar, J., Hess, B., Lindahl, E., and Norlén, L. (2022). Skin permeability prediction with md simulation sampling spatial and alchemical reaction coordinates. Biophysical Journal, 121(20):3837–3849.

[64] Maier, J. A., Martinez, C., Kasavajhala, K., Wickstrom, L., Hauser, K. E., and Simmerling, C. (2015). ff14sb: improving the accuracy of protein side chain and backbone parameters from ff99sb. Journal of chemical theory and computation, 11(8):3696–3713.

[65] Mark, P. and Nilsson, L. (2001). Structure and dynamics of the tip3p, spc, and spc/e water models at 298 k. The Journal of Physical Chemistry A, 105(43):9954–9960.

[66] Markland, T. E. and Ceriotti, M. (2018). Nuclear quantum effects enter the mainstream. Nature Reviews Chemistry, 2(3):0109.

[67] Mobley, D. L., Bannan, C. C., Rizzi, A., Bayly, C. I., Chodera, J. D., Lim, V. T., Lim, N. M., Beauchamp, K. A., Shirts, M. R., Gilson, M. K., et al. (2018a). Open force field consortium: Escaping atom types using direct chemical perception with smirnoff v0. 1. BioRxiv, page 286542.

[68] Mobley, D. L., Bannan, C. C., Rizzi, A., Bayly, C. I., Chodera, J. D., Lim, V. T., Lim, N. M., Beauchamp, K. A., Slochower, D. R., Shirts, M. R., et al. (2018b). Escaping atom types in force fields using direct chemical perception. Journal of chemical theory and computation, 14(11):6076–6092.

[69] Mobley, D. L. and Chodera, J. D. (2023). Machine-learned molecular mechanics force fields from large-scale quantum chemical data. arXiv preprint 2307.07085.

[70] Mobley, D. L., Dumont, É., Chodera, J. D., and Dill, K. A. (2007). Comparison of charge models for fixed-charge force fields: small-molecule hydration free energies in explicit solvent. The Journal of Physical Chemistry B, 111(9):2242–2254.

[71] Mobley, D. L. and Gilson, M. K. (2017). Predicting binding free energies: Frontiers and benchmarks. Annual Review of Biophysics, 46:531–558.

[72] Mobley, D. L. and Guthrie, J. P. (2014). Freesolv: a database of experimental and calculated hydration free energies, with input files. Journal of computer-aided molecular design, 28:711–720.

[73] Naden, L. (2016). Predicting Thermodynamic Properties over Combinatorially Large Chemical Spaces. Phd thesis, School of Engineering and Applied Science, University of Virginia.

[74] Nicholls, A., Mobley, D. L., Guthrie, J. P., Chodera, J. D., Bayly, C. I., Cooper, M. D., and Pande, V. S. (2008). Predicting small-molecule solvation free energies: An informal blind test for computational chemistry. Journal of Medicinal Chemistry, 51(4):769–779.

[75] OpenEye Scientific Software (2024). OpenEye Python Toolkits Documentation. Accessed: 2024-12-18.

[76] Orozco, M., Luque, F., Habibollahzadeh, D., and Gao, J. (1995). The polarization contribution to the free energy of hydration. The Journal of chemical physics, 102(15):6145–6152.

[77] Paszke, A., Gross, S., Chintala, S., Chanan, G., Yang, E., DeVito, Z., Lin, Z., Desmaison, A., Antiga, L., and Lerer, A. (2017). Automatic differentiation in pytorch.

[78] Phillips, J. C., Braun, R., Wang, W., Gumbart, J., Tajkhorshid, E., Villa, E., Chipot, C., Skeel, R. D., Kale, L., and Schulten, K. (2005). Scalable molecular dynamics with namd. Journal of computational chemistry, 26(16):1781–1802.

[79] Plé, T., Adjoua, O., Lagardère, L., and Piquemal, J.-P. (2024). Fennol: an efficient and flexible library for building force-field-enhanced neural network potentials. arXiv preprint 2405.01491.

[80] Prasad, V. K., Otero-de La-Roza, A., and DiLabio, G. A. (2019). Pepconf, a diverse data set of peptide conformational energies. Scientific data, 6(1):1–9.

[81] Ramakrishnan, R., Dral, P. O., Rupp, M., and Von Lilienfeld, O. A. (2014). Quantum chemistry structures and properties of 134 kilo molecules. Scientific data, 1(1):1–7.

[82] Riquelme, M., Lara, A., Mobley, D. L., Verstraelen, T., Matamala, A. R., and Vohringer-Martinez, E. (2018). Hydration free energies in the freesolv database calculated with polarized iterative hirshfeld charges. Journal of chemical information and modeling, 58(9):1779–1797.

[83] Röcken, S., Burnet, A. F., and Zavadlav, J. (2024). Predicting solvation free energies with an implicit solvent machine learning potential. arXiv preprint 2406.00183.

[84] Rodinger, T., Howell, P. L., and Pomès, R. (2005). Absolute free energy calculations by thermodynamic integration in four spatial dimensions. The Journal of chemical physics, 123(3).

[85] Sabanes Zariquiey, F., Galvelis, R., Gallicchio, E., Chodera, J. D., Markland, T. E., and De Fabritiis, G. (2024). Enhancing protein–ligand binding affnity predictions using neural network potentials. Journal of Chemical Information and Modeling, 64(5):1481–1485.

[86] Salahub, D. R., de la Lande, A., Goursot, A., Zhang, R., and Zhang, Y. (2012). Recent progress in density functional methodology for biomolecular modeling. In Challenges and Advances in Computational Chemistry and Physics, volume 13, pages 1–55. Springer.

[87] Schapin, N., Majewski, M., Varela, A., Arroniz, C., and De Fabritiis, G. (2023). Machine learning small molecule properties in drug discovery. arXiv preprint 2308.12354.

[88] Schindler, C. E., Baumann, H., Blum, A., Bose, D., Buchstaller, H.-P., Burgdorf, L., Cappel, D., Chekler, E., Czodrowski, P., Dorsch, D., et al. (2020). Large-scale assessment of binding free energy calculations in active drug discovery projects. Journal of Chemical Information and Modeling, 60(11):5457–5474.

[89] Schnieders, M. J., Baltrusaitis, J., Shi, Y., Chattree, G., Zheng, L., Yang, W., and Ren, P. (2012). The structure, thermodynamics, and solubility of organic crystals from simulation with a polarizable force field. Journal of Chemical Theory and Computation, 8(5):1721–1736.

[90] Scott, W. R., Hünenberger, P. H., Tironi, I. G., Mark, A. E., Billeter, S. R., Fennen, J., Torda, A. E., Huber, T., Krüger, P., and Van Gunsteren, W. F. (1999). The gromos biomolecular simulation program package. The Journal of Physical Chemistry A, 103(19):3596–3607.

[91] Semelak, J. A., Ignacio, P., Huddleston, K. K., Olmos, J., Grassano, J. S., Clemente, C., Drusin, S. I., Marti, M., Lebrero, M. C. G., Roitberg, A. E., et al. (2024). Ani neural networks meet electrostatics: A ml/mm implementation in amber.

[92] Seute, L., Hartmann, E., Stühmer, J., and Gräter, F. (2024). Grappa–a machine learned molecular mechanics force field. arXiv preprint 2404.00050.

[93] Shirts, M. R., Mobley, D. L., and Brown, S. P. (2010). Free-energy calculations in structure-based drug design. Drug Design, 1:61–86.

[94] Shirts, M. R. and Pande, V. S. (2005a). Comparison of efficiency and bias of free energies computed by exponential averaging, the bennett acceptance ratio, and thermodynamic integration. The Journal of chemical physics, 122(14).

[95] Shirts, M. R. and Pande, V. S. (2005b). Solvation free energies of amino acid side chain analogs for common molecular mechanics water models. Journal of Chemical Physics, 122(13):134508.

[96] Singh, S., Gapsys, V., Aldeghi, M., Schaller, D., Rangwala, A. M., White, J. B., Bluck, J. P., Scheen, J., Glass, W. G., Guo, J., et al. (2024). Prospective evaluation of structure-based simulations reveal their ability to predict the impact of kinase mutations on inhibitor binding. bioRxiv, pages 2024–11.

[97] Smith, D. G., Altarawy, D., Burns, L. A., Welborn, M., Naden, L. N., Ward, L., Ellis, S., Pritchard, B. P., and Crawford, T. D. (2021). The molssi qcarchive project: An open-source platform to compute, organize, and share quantum chemistry data. Wiley Interdisciplinary Reviews: Computational Molecular Science, 11(2):e1491.

[98] Smith, J. S., Isayev, O., and Roitberg, A. E. (2017). Ani-1: an extensible neural network potential with dft accuracy at force field computational cost. Chemical science, 8(4):3192–3203.

[99] Swope, W. C., Horn, H. W., and Rice, J. E. (2010). Accounting for polarization cost when using fixed charge force fields. i. method for computing energy. The Journal of Physical Chemistry B, 114(26):8621–8630.

[100] Takaba, K., Friedman, A. J., Cavender, C. E., Behara, P. K., Pulido, I., Henry, M. M., MacDermott-Opeskin, H., Iacovella, C. R., Nagle, A. M., Payne, A. M., et al. (2024). Machine-learned molecular mechanics force fields from large-scale quantum chemical data. Chemical Science, 15(32):12861–12878.

[101] Takaba, K., Pulido, I., Henry, M., MacDermott-Opeskin, H., Chodera, J. D., and Wang, Y. (2023). Espaloma-0.3. 0: Machine-learned molecular mechanics force field for the simulation of protein-ligand systems and beyond. arXiv preprint 2307.07085.

[102] Thaler, S. and Zavadlav, J. (2021). Learning neural network potentials from experimental data via differentiable trajectory reweighting. Nature communications, 12(1):6884.

[103] Tokdar, S. T. and Kass, R. E. (2010). Importance sampling: a review. Wiley Interdisciplinary Reviews: Computational Statistics, 2(1):54–60.

[104] Tse, C. H., Comer, J., Sang Chu, S. K., Wang, Y., and Chipot, C. (2019). Affordable membrane permeability calculations: permeation of short-chain alcohols through pure-lipid bilayers and a mammalian cell membrane. Journal of Chemical Theory and Computation, 15(5):2913–2924.

[105] Turney, J. M., Simmonett, A. C., Parrish, R. M., Hohenstein, E. G., Evangelista, F. A., Fermann, J. T., Mintz, B. J., Burns, L. A., Wilke, J. J., Abrams, M. L., et al. (2012). Psi4: an open-source ab initio electronic structure program. Wiley Interdisciplinary Reviews: Computational Molecular Science, 2(4):556–565.

[106] Unke, O. T., Stöhr, M., Ganscha, S., Unterthiner, T., Maennel, H., Kashubin, S., Ahlin, D., Gastegger, M., Medrano Sandonas, L., Berryman, J. T., et al. (2024). Biomolecular dynamics with machine-learned quantum-mechanical force fields trained on diverse chemical fragments. Science Advances, 10(14):eadn4397.

[107] Van Der Spoel, D., Lindahl, E., Hess, B., Groenhof, G., Mark, A. E., and Berendsen, H. J. (2005). Gromacs: fast, flexible, and free. Journal of computational chemistry, 26(16):1701–1718.

[108] Wang, L., Behara, P. K., Thompson, M. W., Gokey, T., Wang, Y., Wagner, J. R., Cole, D. J., Gilson, M. K., Shirts, M. R., and Mobley, D. L. (2024a). The open force field initiative: Open software and open science for molecular modeling. The Journal of Physical Chemistry B, 128(29):7043–7067.

[109] Wang, M., Zheng, D., Ye, Z., Gan, Q., Li, M., Song, X., Zhou, J., Ma, C., Yu, L., Gai, Y., et al. (2019). Deep graph library: A graph-centric, highly-performant package for graph neural networks. arXiv preprint 1909.01315.

[110] Wang, Y. (2023). Graph Machine Learning for (Bio) Molecular Modeling and Force Field Construction. PhD thesis, Weill Medical College of Cornell University.

[111] Wang, Y., Fass, J., Kaminow, B., Herr, J. E., Rufa, D., Zhang, I., Pulido, I., Henry, M., Macdonald, H. E. B., Takaba, K., et al. (2022). End-to-end differentiable construction of molecular mechanics force fields. Chemical Science, 13(41):12016– 12033.

[112] Wang, Y., Pulido, I., Takaba, K., Kaminow, B., Scheen, J., Wang, L., and Chodera, J. D. (2024b). Espalomacharge: Machine learning-enabled ultrafast partial charge assignment. The Journal of Physical Chemistry A, 128(20):4160–4167.

[113] Wong, T.-T. (2015). Performance evaluation of classification algorithms by k-fold and leave-one-out cross validation. Pattern recognition, 48(9):2839–2846.

[114] Wu, D. and Kofke, D. A. (2005). Phase-space overlap measures. i. fail-safe bias detection in free energies calculated by molecular simulation. The Journal of chemical physics, 123(5).

[115] Wu, Z., Ramsundar, B., Feinberg, E. N., Gomes, J., Geniesse, C., Pappu, A. S., Leswing, K., and Pande, V. (2018). Moleculenet: a benchmark for molecular machine learning. Chemical science, 9(2):513–530.

[116] Zaki, M. J. and Hsiao, C.-J. (2002). Charm: An efficient algorithm for closed itemset mining. In Proceedings of the 2002 SIAM international conference on data mining, pages 457–473. SIAM.

[117] Zhao, Y. and contributors (2024). timemachine: A high-performance differentiable molecular dynamics and force-field engine. https://github.com/proteneer/timemachine. Accessed: 2025-12-09.

[118] Zwanzig, R. W. (1954). High-temperature equation of state by a perturbation method. i. nonpolar gases. The Journal of Chemical Physics, 22(8):1420–1426.

